# Single-cell functional genomics of natural killer cell evasion in blood cancers

**DOI:** 10.1101/2022.08.22.504722

**Authors:** Olli Dufva, Sara Gandolfi, Jani Huuhtanen, Olga Dashevsky, Khalid Saeed, Jay Klievink, Petra Nygren, Jonas Bouhlal, Jenni Lahtela, Anna Näätänen, Bishwa R Ghimire, Tiina Hannunen, Pekka Ellonen, Hanna Duàn, Jason Theodoropoulos, Essi Laajala, Jouni Härkönen, Petri Pölönen, Merja Heinäniemi, Shizuka Yamano, Ryosuke Shirasaki, David Barbie, Jennifer Roth, Rizwan Romee, Michal Sheffer, Harri Lähdesmäki, Dean A. Lee, Ricardo De Matos Simoes, Matti Kankainen, Constantine S Mitsiades, Satu Mustjoki

## Abstract

Natural killer (NK) cells are emerging as a promising therapeutic option in cancer. To better understand how cancer cells evade NK cells, we studied interacting NK and blood cancer cells using single-cell and genome-scale functional genomics screens. At single-cell resolution, interaction of NK and cancer cells induced distinct activation states in both cell types depending on the cancer cell lineage and molecular phenotype, ranging from more sensitive myeloid to more resistant B-lymphoid cancers. CRISPR screens uncovered cancer cell-intrinsic genes driving sensitivity and resistance, including antigen presentation and death receptor signaling mediators, adhesion molecules, protein fucosylation genes, and transcriptional regulators. CRISPR screens with a single-cell transcriptomic readout revealed how these cancer cell genes influenced the gene expression landscape of both cell types, including regulation of activation states in both cancer and NK cells by IFNγ signaling. Our findings provide a resource for rational design of NK cell-based therapies in blood cancers.

**HIGHLIGHTS:**

- Transcriptomic states of interacting NK cells and cancer cells depend on cancer cell lineage
- Molecular correlates of increased sensitivity of myeloid compared to B-lymphoid cancers include activating receptor ligands NCR3LG1, PVR, and ULBP1
- New regulators of NK cell resistance from 12 genome-scale CRISPR screens include blood cancer-specific regulators SELPLG, SPN, and MYB
- Single-cell transcriptomics CRISPR screens targeting 65 genome-wide screen hits identify MHC-I, IFNy, and NF-κB regulation as underlying mechanisms

## INTRODUCTION

NK cells are cytotoxic innate lymphoid cells which can directly eliminate cancer cells through secretion of cytolytic granules and trigger an immune response via secretion of immunomodulatory cytokines (Chiossone et al., 2018). NK cell activation relies on a balance between activating and inhibitory signals derived from surface receptors engaged with cognate ligands on target cells (Lanier, 2003). Data on NK cells driving the graft-versus-leukemia effect in allogeneic hematopoietic stem cell transplantation (Ruggeri et al., 2002) and more recently on the efficacy of chimeric antigen receptor (CAR) NK cells (Liu et al., 2020) have provided encouraging evidence on the therapeutic potential of NK cells in blood cancers. As a result, NK cell-based immunotherapies (adoptive transfer of allogeneic NK cells, bispecific NK engager antibodies, and CAR NK cells) are actively pursued in patients with hematological malignancies (Myers and Miller, 2021; Shimasaki et al., 2020).

Single-cell analyses have provided unbiased transcriptional profiles of immune cell subsets derived from healthy and malignant tissues, as well as of transcriptional programs in tumor cells (Cheng et al., 2021; Crinier et al., 2018; Jerby-Arnon et al., 2018; Pfefferle et al., 2019; Smith et al., 2020; Yang et al., 2019; Zheng et al., 2021). However, these approaches are not optimally positioned to capture the dynamic changes in tumor and immune cells when these cells interact. How cytotoxic lymphocytes such as NK cells react to tumor cell challenge and how tumor cells in turn respond by altering their transcriptional states has not been investigated in a systematic, transcriptome-wide manner.

Genome-scale CRISPR screens have revealed cancer cell-intrinsic mechanisms of evasion from T cell killing in solid tumors (Kearney et al., 2018; Lawson et al., 2020; Patel et al., 2017) and hematological malignancies (Dufva et al., 2020a; Singh et al., 2020), confirming that unbiased functional genomics can reveal underappreciated aspects of immune-cancer cell interactions. CRISPR screens and measurement of NK cell sensitivity across a large panel of genotypically diverse cell lines using PRISM (Yu et al., 2016) identified molecular factors regulating sensitivity of solid tumor cells to NK cell mediated killing (Sheffer et al., 2021). In contrast, a systematic evaluation of resistance and sensitivity to NK cell therapies across hematological malignancies has not been performed, limiting our biological understanding of endogenous NK cell-mediated anti-cancer immunity as well as the therapeutic use of NK cells for blood cancers, the main context for which clinical proof-of-concept is available and rapid further development is anticipated for NK cell-based therapies.

As a result, several key questions in translating the potential of NK cells as effective therapies in blood cancers remain unanswered. How do NK cells and cancer cells respond to their interaction by changing their transcriptional profiles and do these changes differ depending on the phenotype or genetic makeup of the cancer cells? Are there previously undiscovered mechanisms mediating NK cell cytotoxicity, including mechanisms unique to blood cancers? Do genetic or epigenetic alterations in blood cancer cells from individual patients influence NK cell sensitivity/resistance mechanisms, ultimately leading to differences in sensitivity to NK cells between molecular subtypes of blood cancers? Linking genomic subtypes of blood cancers to NK cell sensitivity would enable identification of patient groups more likely to benefit from NK cell-based immunotherapy, and pinpoint molecular mechanisms that could be therapeutically targeted to sensitize resistant malignancies to NK-cell based therapies.

Here, we sought to answer these questions by combining multiplexed single-cell RNA-seq (scRNA-seq) profiling of interacting NK cells and cancer cells, PRISM-based profiling of NK cell sensitivity across a panel of blood cancer cell lines, and genome-scale and single-cell transcriptomic CRISPR screens of cancer-cell intrinsic NK cell resistance mechanisms (Figure 1A). By integrating these data and patient genomic profiles, we provide a comprehensive landscape of functionally validated genetic mechanisms which influence how NK cells recognize and kill malignant hematopoietic cells. The results offer a roadmap to facilitate development of NK-cell based immunotherapy for blood cancers and beyond. The data are available for interactive exploration at https://immunogenomics.shinyapps.io/nkheme/.

**Figure 1.**
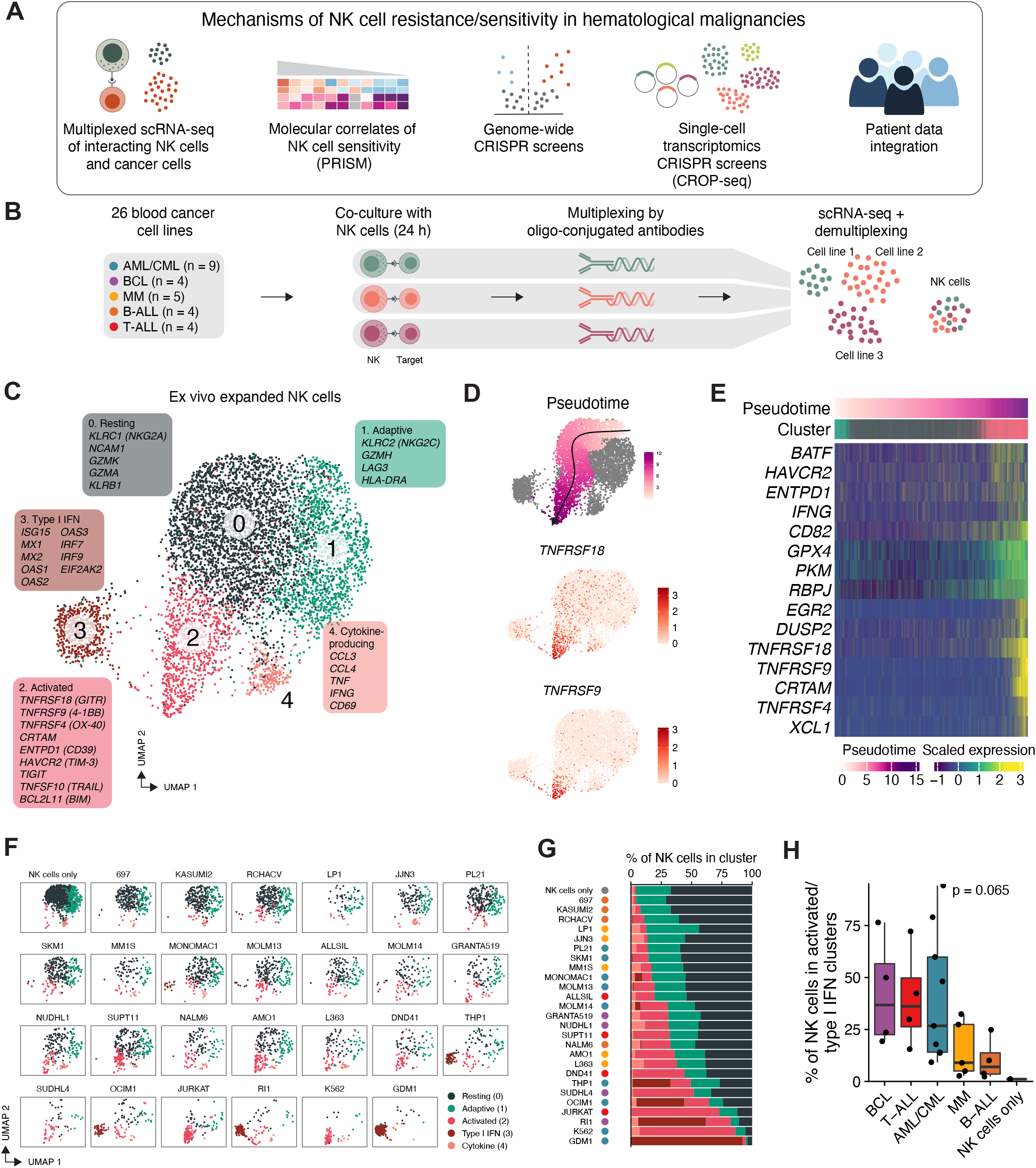
Single-cell transcriptomics of NK cells interacting with blood cancer cells. (A) Overview of the study. (B) Multiplexed co-culture scRNA-seq workflow. (C) UMAP visualization of expanded NK cells from all conditions, including co-culture with 26 cell lines and NK cells cultured alone. Cells are colored based on the clusters and marker genes are shown for each cluster selected from the genes significantly overexpressed in each cluster compared to other clusters. (D) UMAP visualizations as in C showing pseudotime starting from cluster 0 (resting) and ending in cluster 2 (activated) (top), and expression of genes enriched in the activated cluster (middle and bottom). Arrow indicates the pseudotime trajectory. (E) Heatmap of selected genes upregulated in the activated cluster. Cells included in the trajectory in D are shown ordered by pseudotime. (F) UMAP visualizations of expanded NK cells across all conditions, including co-culture with 26 cell lines and NK cells cultured alone. (G) Bar plot of percentages of NK cells belonging to different clusters in each condition, including co-culture with 26 cell lines and NK cells cultured alone. Conditions are ordered by the combined percentage of cells in resting and adaptive (clusters 0 and 1) in decreasing order. (H) Box plot of percentages of NK cells belonging to different clusters in each condition stratified by cancer type. P value between all groups is obtained using a Kruskal-Wallis test. Boxes indicate IQR with a line at the median. Whiskers represent the min and max values at most 1.5 IQR from the quartiles. See also Figure S1 and Table S1.

## RESULTS

### Multiplexed scRNA-seq defines phenotypic changes in NK cells interacting with blood cancer cells

Defining how NK cells and cancer cells change their phenotype upon their interaction is essential for understanding potential mechanisms of resistance. To comprehensively profile the cell states of interacting NK cells and blood cancer cells, we cultured 26 different cell lines representing diverse hematologic neoplasms either alone or with NK cells derived from a single donor (Figure 1B, Table S1). The cancer types included acute and chronic myeloid leukemia (AML and CML), B and T cell acute lymphoblastic leukemia (B-ALL and T-ALL), B cell lymphoma (BCL), and multiple myeloma (MM). We studied both NK cells extracted directly from the peripheral blood and NK cells expanded *ex vivo* using feeder cells and IL-2, corresponding to those used in adoptive NK cell immunotherapy trials (Ciurea et al., 2017; Liu et al., 2020). After 24 h co-culture, we labeled the cells from each monoculture or co-culture condition with oligonucleotide-conjugated antibodies against ubiquitously expressed surface proteins (with different oligonucleotide for each mono- or co-culture), enabling multiplexing in the scRNA seq using the cell hashing method (Stoeckius et al., 2018). Across the 82 scRNA-seq samples, we obtained a total of 61,715 cells (753 cells per sample on average) classified as singlets based on hashtag barcodes, including 8,851 NK cells (182 cells per sample on average) and 52,864 target cells (645 cells per sample on average) (Figure S1A-B; Table S1).

We first focused on NK cells. After correcting for cell cycle and batch effects, unsupervised clustering of the *ex vivo* expanded NK cells from all conditions revealed five distinct clusters (see Methods; Figures 1C and S1C). Most monocultured NK cells belonged to two clusters: resting NK cells (cluster 0) expressing markers of CD56bright NK cells (*NCAM1/CD56, KLRC1*/*NKG2A, GZMK*) (Crinier et al., 2018; Pfefferle et al., 2019; Smith et al., 2020; Yang et al., 2019), consistent with the CD56bright phenotype of expanded NK cells (Denman et al., 2012; Lieberman et al., 2018), and adaptive NK cells (cluster 1) based on expression of *KLRC2*/*NKG2C* and *LAG3* (Holmes et al., 2021; Merino et al., 2019) (Figure 1C). In contrast, the remaining three NK cell clusters (clusters 2, 3, and 4) were present in very small quantities when NK cells were cultured alone, but were enriched in the co-culture conditions with tumor cells. NK cells with an activated phenotype (cluster 2) expressed genes encoding several co-stimulatory receptors *TNFRSF18*/*GITR, TNFRSF9*/*4-1BB, TNFRSF4*/*OX-40*, and *CRTAM*, inhibitory receptors such as *HAVRC2*/*TIM-3* and *TIGIT*, immediate-early genes *DUSP2, DUSP4*, and *EGR2*, the death receptor ligand *TNFSF10*/*TRAIL*, and *ENTPD1/CD39* associated with tumor-reactive T cells and exhaustion (Gupta et al., 2015; Sade-Feldman et al., 2018; Simoni et al., 2018). On a pseudotemporal trajectory from the resting to the activated cluster, *BATF, HAVCR2*, and *ENTPD1* were expressed early in the transition to the activated state, whereas *TNFRSF18, TNFRSF9, TNFRSF4, and CRTAM* marked the terminal point of the NK cell activation state spectrum (Figures 1D-1E). Other clusters enriched upon target cell enrichment included NK cells with high type I interferon (IFN) signature (cluster 3) expressing antiviral genes such as *MX1, MX2, OAS1, OAS2*, and *OAS3* and cytokine-producing NK cells expressing several cytokine genes including *CCL3, CCL4, TNF*, and *IFNG* (cluster 4).

Different cell lines induced distinct changes in the phenotype of NK cells. Some cell lines such as K562 (CML), JURKAT (T-ALL), and SUDHL4 (BCL) induced over 50% of NK cells to transition into the activated state (cluster 2), compared to less than 20% with several B-ALL and MM cell lines (Figures 1F-1H). In contrast to the gradual differences in transition to the activated cluster 2 across cell lines, the type I IFN NK cell state (cluster 3) was only induced by certain cell lines, including the AML lines GDM1, OCIM1, and THP1, and the BCL line RI1. As almost all cell lines were matched with the NK cells with regard to HLA-C1/HLA-C2 groups (Table S1), cancer cell features other than human leukocyte antigen (HLA) types are likely responsible for the distinct NK cell activation states in the co-culture conditions.

NK cells extracted directly from peripheral blood mononuclear cells (PBMC) from the same donor without expansion showed largely similar responses to co-culture with cancer cells, indicating that the observed activation states are also relevant to NK cells normally found in circulation (Figures S1D-S1G). In PBMC NK cells, however, the transition to the activated cluster was tightly coupled with the cytokine cluster (Figure S1G), in contrast to expanded NK cells where the cytokine cluster varied independently of the activated cluster. Together, these findings indicate that NK cells shift into activated states with an altered repertoire of co-stimulatory and co-inhibitory receptors in response to engagement with cancer cells, and that the magnitude and direction of the transition varies depending on the target cells.

### Transcriptomic responses of blood cancer cells to NK cell attack

Having defined the changes in NK cell phenotypes resulting from co-culture, we examined the transcriptomic responses induced in cancer cells by the NK cell attack. Comparison of all expanded NK cell-treated target cells with the untreated controls indicated a strong interferon γ (IFNγ) response induced by NK cell treatment (Figure 2A; Table S1). A core set of 17 genes significantly induced in over 75% of the cell lines comprised the class I major histocompatibility complex (MHC-I) genes (*B2M, HLA-A, HLA-B, HLA-C*, and *HLA-E*), JAK-STAT signaling genes (*STAT1, IRF1, IRF9*), immunoproteasome genes (*PSMB8, PSMB9, PSBM10, PSME1, PSME2*), the ubiquitin-conjugating enzyme gene *UBE2L6*, granulysin (*GNLY)*, and the chemokine-ligand 5 gene (*CCL5)* (Figure 2B). To test which cell lines showed the strongest transcriptomic responses, we ranked the cell lines by the log2 fold change (log2FC) of a score comprising the 17 core NK cell-induced genes (Figures 2B and S1H-S1I). This revealed particularly strong responses in T-ALL and myeloid AML and CML cell lines compared to other blood cancer types (Figure 2C). Transcriptomic changes induced by non-expanded PBMC-derived NK cells in blood cancer target cells were similar but less pronounced than those observed with expanded NK cells (Figure S1I).

**Figure 2.**
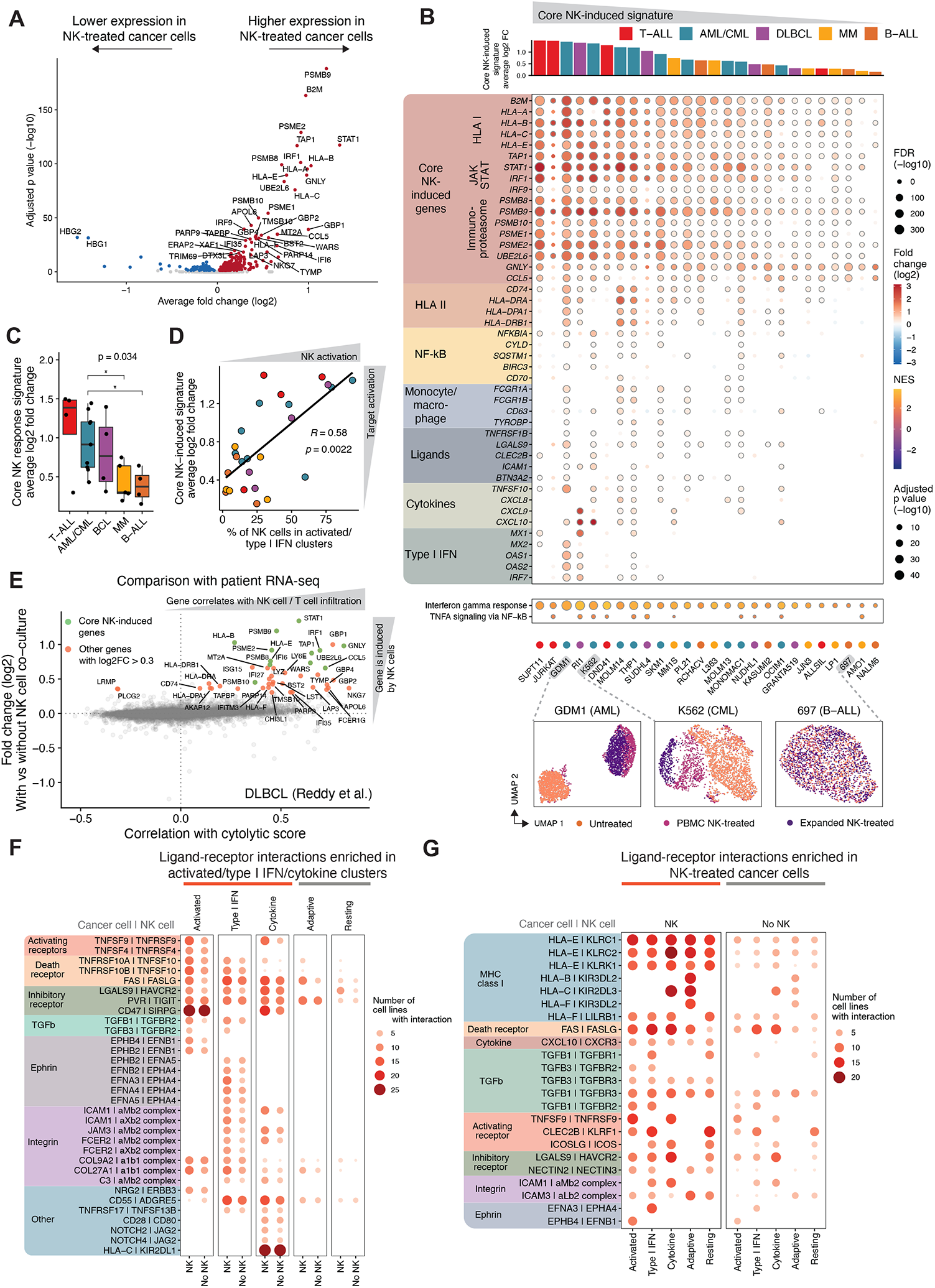
Transcriptomic responses of blood cancer cells to NK cell attack. (A) Volcano plot of differentially expressed genes between all NK-cell treated cancer cell lines and the same cell lines cultured alone. Red dots indicate genes significantly (FDR < 0.05) enriched in NK-treated cells and blue dots indicate genes enriched in untreated cells. (B) Dot plot of selected genes upregulated in cancer cells upon exposure to expanded NK cells. The genes shown include the core NK cell-induced genes (17 genes induced in at least 75% of the cell lines) and other genes induced in subsets of the cell lines grouped based on functional categories. The cell lines are ordered based on the log2 fold change of a score comprising the core NK cell-induced genes shown as a bar plot at the top. Color indicates log2 fold change between conditions and dot size indicates the negative log10 FDR. Only dots where P < 0.05 are shown, and circled dots indicate FDR < 0.05. At the bottom, examples of UMAP visualizations of the cell lines GDM1, K562, and 697 are shown colored according to the co-culture condition. (C) Box plot of the log2 fold change of the core NK cell-induced gene score between NK-treated and untreated cells stratified by cancer type. P value between all groups is obtained using a Kruskal-Wallis test and p values between each pair using Wilcoxon rank sum tests followed by Benjamini-Hochberg (BH) adjustment. Only p values for significant pairs are shown (* < 0.05). Boxes indicate IQR with a line at the median. Whiskers represent the min and max values at most 1.5 IQR from the quartiles. (D) Scatter plot comparing NK cell activation (percentage of NK cells in activated and type I IFN clusters after co-culture with the cell lines) and core NK cell response gene set induction in NK-treated target cells (log2 fold change in core NK cell response score compared to untreated). Correlation coefficient and p value are obtained using Spearman’s rank correlation. Dots are colored according to cancer type as in C. (E) Scatter plot comparing genes induced by NK cell co-culture in target cells and genes correlating with NK and T cell infiltration (cytolytic score) in DLBCL patient samples from Reddy et al. Genes with significant correlation and differential expression (FDR < 0.05) and scRNA-seq log2 fold change > 0.3 are labeled in red. Genes included in the core NK-induced genes are labeled in green. (F) Dot plot of ligand-receptor interactions induced by NK cell activation. Dot size and color indicates the number of cell lines (out of total 26) having significant interactions with each cluster of expanded NK cells according to CellPhoneDB. Shown are interactions most enriched in the three activation-related clusters (activated, type I IFN, cytokine-producing) compared to the two other clusters (resting and adaptive), indicating interactions induced by NK cell activation states. Interactions are ordered based on functional categories. (G) Dot plot of ligand-receptor interactions induced by cancer cell response to NK cells. Dot size and color indicates the number of cell lines (out of total 26) having significant interactions with each cluster of expanded NK cells according to CellPhoneDB. Shown are interactions most enriched in the NK cell-treated conditions compared to untreated, indicating interactions induced by cancer cell response to NK cell attack. Interactions are ordered based on functional categories. See also Figure S1 and Table S1.

To uncover distinct transcriptional programs induced in subsets of the cell lines, we examined 200 genes most variably induced across cell lines by unsupervised clustering (Figure S1H). This highlighted IFNγ-associated programs, including the core set of NK-induced genes as well as cytokines such as *CXCL9, CXCL10*, and *TNFSF10* (*TRAIL*). MHC class II (MHC-II) genes were induced more prominently in monocytic cells and several B cell lines. A program related to differentiation of myeloid cells towards monocytes, macrophages, and neutrophils was induced in AML cells, including genes such as *FCGR1A, CD63*, cathepsins, and the TIM-3 ligand *LGALS9*, suggesting that NK cell attack may induce maturation of monocytic leukemias, as has been reported in response to IFNγ (Matsuo et al., 1997). Strikingly, a type I IFN signature was induced in many of the same cells which induced a similar signature in the NK cells (GDM1, R1, and THP1), indicating a coordinated response in both cell types evoked by their interaction (Figures 2B and S1H). When comparing the core NK signature induced in the target cells with the percentage of NK cells shifting towards the activated/type I IFN phenotypes, we indeed observed a positive correlation between NK cell and target cell activation (Figure 2D).

To determine whether similar NK cell-induced signatures in cancer cells could be found in patient data, we compared the differentially expressed genes with vs. without NK cell co-culture *in vitro* to genes correlating with infiltration by cytotoxic lymphocytes (NK or T cells) in patients with diffuse large B cell lymphoma (DLBCL) (Reddy et al., 2017) or AML (2013) (Figures 2E and S1J). The majority of the NK-induced genes, particularly the 17-gene core set, correlated positively with NK cell infiltration, suggesting that similar transcriptomic responses occur *in vivo*.

Given the observed changes in expression of various receptors in NK cells and ligands in cancer cells, we used the CellPhoneDB (Efremova et al., 2020) to study which ligand-receptor interactions would be unique to cells in the interacting states, as opposed to resting states. Several interactions were more frequent between the cancer cells and co-culture-related NK cell clusters (activated, type I IFN, or cytokine), compared to the resting or adaptive-like clusters (Figure 2F). These included ligands for activating and inhibitory receptors (*TNFSF9*-*TNFRSF9, TNFSF4*-*TNFRSF4, LGALS9*-*HAVCR2*), death receptors (*TNFRSF10A/B*-*TNFSF10, FAS*-*FASLG*), cytokines (*TGFB1/3*-*TGFBR2*), and adhesion molecules (ICAM1-aMb2/aXb2 complex, EFNA3/4/5-EPHA4), indicating that new interactions emerge upon NK cell transition into the cancer cell-induced states. In turn, several interactions were induced by transition of the cancer cells to the NK cell-treated states, including interactions of MHC-I with inhibitory receptors, and the engagement of the IFNγ-regulated cytokine CXCL10 with the CXCR3 receptor (Figure 2G). Taken together, the transcriptomic responses of tumor cells to NK cell attack depend on the lineage and correlate with NK cell activation, resulting in new ligand-receptor interactions not found in the resting state.

### Molecular correlates of NK cell sensitivity across blood cancers

To study cancer cell sensitivity to NK cells across different lineages and stages of maturation, we performed phenotypic studies on a pool of 63 molecularly-annotated DNA-barcoded blood cancer cell lines, including myeloid and lymphoblastic leukemia, DLBCL, and MM cells (PRISM system, Figure 3A) (Yu et al., 2016). We quantified the dose-dependent responses to primary NK cells using the relative abundance of barcodes in treated cells compared to controls (Table S2), followed by integrated computational analyses to identify candidate molecular markers correlating with tumor cell sensitivity or resistance to NK cells.

**Figure 3.**
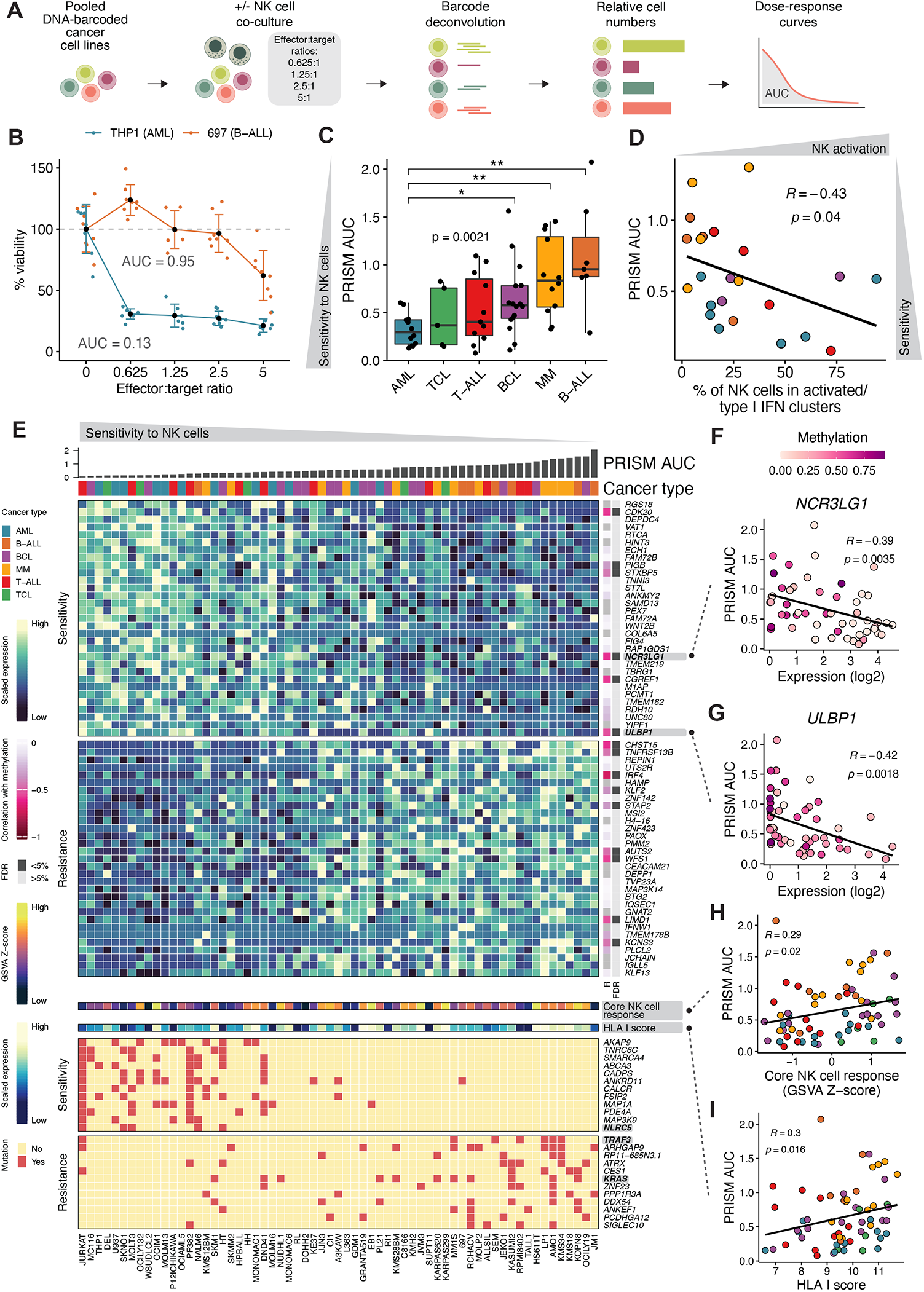
PRISM screen of NK cell sensitivity across blood cancer cell lines. (A) PRISM screen workflow. (B) Examples of dose-response curves of NK cell sensitivity at various effector-to-target ratios obtained using PRISM for two cell lines, 697 (B-ALL) and THP-1 (AML). The percent viability of the target cells at each effector-to-target ratio is obtained by normalizing the luminescence values to the mean of the untreated (0:1) condition. Colored dots represent technical replicates (n = 8), black dots indicate mean, error bars indicate standard deviation, and the area under the dose-response curve (AUC) is shown next to the curve. Lower AUC denotes sensitivity to NK cells and higher AUC resistance. (C) Box plot of the sensitivity of different hematological malignancies to NK cell cytotoxicity shown as AUC of the dose-response curve. P value between all groups shown in the figure is obtained using a Kruskal-Wallis test and p values between each pair using Wilcoxon rank sum tests followed by Benjamini-Hochberg (BH) adjustment. Only p values for significant pairs are shown (* < 0.05, ** < 0.01). Boxes indicate IQR with a line at the median. Whiskers represent the min and max values at most 1.5 IQR from the quartiles. (D) Scatter plot comparing NK cell activation (percentage of NK cells in activated and type I IFN clusters after co-culture with the cell lines) and sensitivity of the cell lines to NK cells in PRISM quantified as AUC of the dose-response curve. Correlation coefficient and p value are obtained using Spearman’s rank correlation. Dots are colored according to cancer type as in C. (E) Heatmap of molecular correlates of sensitivity to NK cells across blood cancer cell lines. Cell lines are ordered by decreasing sensitivity (increasing PRISM AUC). Expression of genes and mutations most highly correlated with sensitivity or resistance to NK cells are shown. For the shown genes, correlation of expression with methylation is shown on the right, with a separate column indicating significance of the correlation at 5% FDR. In addition, expression of the core NK cell response gene set derived from co-culture scRNA-seq experiments as GSVA score and HLA I score summarizing expression of MHC-I complex genes are shown. (F) Scatter plot comparing expression of *NCR3LG1* with NK cell sensitivity (PRISM AUC). Dot color indicates *NCR3LG1* methylation. Correlation coefficient and p value are obtained using Spearman’s rank correlation. (G) Scatter plot comparing expression of *ULBP1* with NK cell sensitivity (PRISM AUC). Dot color indicates *ULBP1* methylation. Correlation coefficient and p value are obtained using Spearman’s rank correlation. (H) Scatter plot comparing expression of the core NK cell response gene set as GSVA score with NK cell sensitivity (PRISM AUC). Correlation coefficient and p value are obtained using Spearman’s rank correlation. (I) Scatter plot comparing HLA I score summarizing expression of HLA I complex genes with NK cell sensitivity (PRISM AUC). Correlation coefficient and p value are obtained using Spearman’s rank correlation. See also Figure S2 and Table S2.

We observed substantial heterogeneity across blood cancer types in their sensitivity to NK cells (Figures 3B-3C and S2A). AML cell lines were the most sensitive, whereas B-ALL cells were generally resistant. MM cell lines were on average relatively resistant and T-ALL cell lines were sensitive, but both showed a high degree of variation between individual cell lines, implying the existence of sensitive and resistant subgroups. Such heterogeneity could also be observed when comparing the sensitivity of each individual cell line and the corresponding percentage of NK cells shifting towards the activated/type I IFN clusters (Figures 3D, 1H, and S2B). Not all responsive cell lines elicited a strong transition of the effector cells into an activated/type I IFN phenotype. However, all of the resistant cell lines we tested failed to induce this phenotype, indicating that the ability to induce an NK cell activation state is one of the possible mechanisms explaining the heterogeneity across disease subtypes (Figure 1G).

We next asked which molecular features could explain the observed variation in NK cell sensitivity. We explored correlations of the AUC of NK cell sensitivity to gene expression, DNA methylation, mutations, microRNAs, proteomics, and metabolomics in the Cancer Cell Line Encyclopedia (CCLE) multi-omics data (Barretina et al., 2012; Ghandi et al., 2019) (Figures 3E and S2C). Among the genes most highly correlated with sensitivity to NK cells were the activating receptor ligands *NCR3LG1* and *ULBP1* (false discovery rate (FDR) cutoff 15%). Expression of *NCR3LG1* and *ULBP1* correlated negatively with methylation of their promoter regions (Figures 3F-3G), suggesting methylation-based epigenetic regulation as a basis for cancer type heterogeneity of these activating signals. Genes that correlated with resistance to NK cells included components of the alternative NF-κB pathway (*TNFRSF13B/TACI, MAP3K14/NIK*).

In gene set variation analysis (GSVA), the core NK cell response signature derived from single-cell data (Figure 2B) correlated with resistance (Figure 3H) concordantly with the class I HLA expression score (Figure 3I). This observation supports the hypothesis that pre-existing expression of those adaptive response molecules may contribute to the intrinsic resistance of cancer cells to NK cells. Although not meeting the criteria for significance after multiple testing correction, several genetic alterations and protein, miRNA, and metabolite levels were associated with sensitivity to NK cells (Figures 3E and S2C). *TRAF3* mutations mostly found in MM and *KRAS* mutations found in multiple cancer types correlated with resistance (Figure 3E). In contrast, *NLRC5* mutations were associated with sensitivity, consistent with the MHC-I regulatory function of NLRC5 (Meissner et al., 2010). These molecular correlates may partially reflect different patterns across cancer types, which itself emerged as a key determinant of NK cell sensitivity, but also potential mechanistic roles of the respective molecules in regulating NK cell response.

### CRISPR screens identify genetic determinants of NK cell sensitivity and resistance in hematological malignancies

To further explore mechanisms that could potentially explain the observed heterogeneity in response to NK cells, we performed genome-scale CRISPR screens in cell lines representing a spectrum of hematologic malignancies, with variable baseline sensitivity to NK cells. We transduced cell lines from BCL (SUDHL4), B-ALL (NALM6), MM (MM1.S, LP1, KMS11), CML (K562), and AML (MOLM14) with either the GeCKO v2 library (Sanjana et al., 2014) or the Brunello library (Doench et al., 2016) for loss-of-function (LOF) screens (see Methods). MM cell lines were also transduced with the gain-of-function (GOF) Calabrese library (Sanson et al., 2018) in order to uncover genes with low expression at baseline whose overexpression might alter the cancer cell response to NK cells. Pools of transduced tumor cells were then co-cultured with IL-2-expanded donor-derived NK cells or the NK cell line KHYG1 for a period ranging from a minimum of 24 hours to up to two weeks (Figure 4A; Table S2).

**Figure 4.**
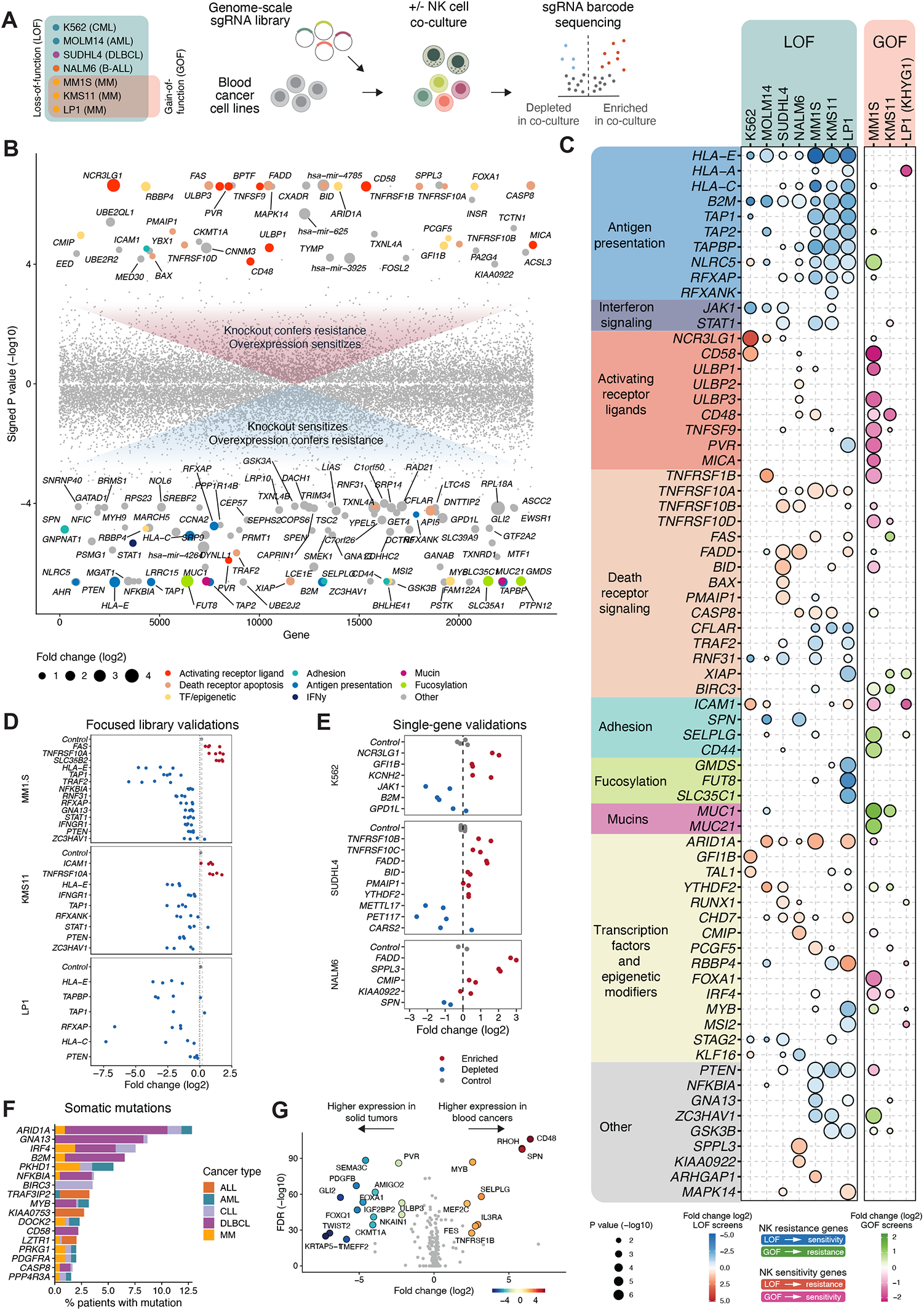
Genome-scale CRISPR screens of NK cell resistance and sensitivity in hematological malignancies. (A) Genome-scale CRISPR screen workflow. (B) Scatter plot of genes conferring resistance or sensitivity to NK cells in genome-scale CRISPR screens. The y axis indicates the p value multiplied by the sign of the log2 fold change. Genes with p < 0.0001 and absolute value of the log2 fold change > 0.75 in at least one screen are shown for the screen with the highest significance for each gene. Dot size indicates the absolute value of the log2 fold change of the labeled genes. Genes are in a randomly sampled order on the x axis. Selected genes out of those with p < 0.0001 included in Figure 1C are colored based on the functional categories. (C) Dot plot of genes conferring resistance or sensitivity to NK cells in genome-scale CRISPR screens. Shown are selected genes out of those with p < 0.0001 in at least one screen or genes validated in separate assays. Color indicates log2 fold change between NK cell-treated and untreated conditions and dot size indicates the negative log10 p value, with only dots where p < 0.05 shown. In LOF screens, blue color indicates depletion of the edited cells and red color enrichment with NK cell treatment. In GOF screens, green color indicates enrichment and pink indicates depletion of the edited cells with NK cell treatment. (D) Selected genes validated using pooled CRISPR screens with focused libraries targeting genome-wide screen hits in MM cell lines. Plots show log2 fold change between NK-treated and untreated cells for each guide RNA. Olfactory receptor (OR) genes were used as a control and shown in gray is the average of all OR genes sgRNAs log2 fold change, while the dashed lines represent the 95% confidence interval. (E) Genes validated using single gene targeting of genome-wide screen hits with a luciferase cytotoxicity assay. Plots show log2 fold change of luciferase-based cell viability between NK-treated and untreated cells for each guide RNA. Dots indicate means of technical replicates for each sgRNA, with two sgRNAs used for each gene. (F) Frequency of mutations in CRISPR screen hit genes in patients with hematological malignancies. Stacked bars indicate percentages of mutated samples of each cancer type cohort. Only genes with mutations in > 1.5% patients cumulatively in all cancer types are shown. (G) Volcano plot comparing expression of CRISPR screen hit genes between blood cancer and solid tumor cell lines in CCLE RNA-seq data. Adjusted p values (-log10) on the y axis are obtained using Wilcoxon rank sum tests followed by Bonferroni correction and log2 fold change of medians is shown on the x axis. Genes with log2 fold change > 2 or < −2 and negative log2 adjusted p value > 20 are labeled and shown as colored dots. See also Figures S4 and S5 and Tables S3 and S4.

As expected, LOF of molecules belonging to the MHC-I complex and the antigen presentation machinery were significantly depleted across all cell lines, reflecting the missing-self mechanism of NK cell activation and thus providing an internal quality control of the screens (Figures 4B-4C and S3A). In addition, other depleted knockouts included genes controlling MHC-I transcriptional regulation, such as *NLRC5, RFXAP*, and *RFXANK*, as well as peptide-loading complex components *TAP1, TAP2*, and *TAPBP*, indispensable for MHC-I surface expression. Also, LOF of genes belonging to the IFNγ signaling pathway – namely *JAK1* and *STAT1* – were significantly depleted in co-culture conditions compared to controls across most cell lines, in keeping with the role of IFNγ-related NK cell response signature in driving reduced NK cell sensitivity observed in the PRISM data (Figures 3H-3I). These findings highlight that genes associated with IFNγ signaling and the antigen presentation machinery are prominent suppressors of NK cell killing regardless of the lineage and the disease subtype.

In contrast, LOF of the death receptor signaling pathway genes – specifically *TNFRSF1B, TNFRSF10A, TNFRSF10B, FAS, FADD, BID*, and *CASP8* – was associated with decreased response to NK cell killing, while the overexpression of *TNFRSF10D* conferred higher sensitivity in two MM cell lines (Figure 4B-C). Consistent with these findings, LOF of the negative regulators of death receptor signaling *CFLAR* (c-FLIP), *TRAF2*, and *XIAP* sensitized multiple cell lines to NK cells. An integrative analysis of data obtained in NALM6 cells treated with either NK cells or CAR T cells (Dufva et al., 2020a) indicated *FADD* and *TNFRSF10B* LOF as a shared mechanism of resistance, consistent with death receptor signaling mediating both NK and T cell cytotoxicity (Figure S3B). Gene set enrichment analysis (GSEA) revealed enrichment of pathways including FASL/CD95L signaling, transcription of death receptors and ligands, TRAIL signaling and TNFR1 induced NFkB signaling pathway as well as IFNγ signaling, antigen processing and presentation, class I HLA assembly and peptide loading in multiple cell lines, supporting the broad relevance of death receptor apoptosis and MHC-I in regulating NK cell cytotoxicity (Figure S3C).

The overexpression of several activating NK cell receptor ligands sensitized cells to NK cell-mediated killing in our MM1.S GOF screen. These included the CD2 ligand *CD58*, the NKG2D ligands *ULBP1, ULBP2, ULBP3*, and *MICA*, the 2B4 ligand *CD48*, the DNAM-1 ligand *PVR*, as well as the TNFSF9, the ligand for the 4-1BB receptor induced in NK cells by tumor challenge in our scRNA-seq analyses. However, the activating receptor ligands showed a heterogeneous pattern across cell lines in the LOF screens. *NCR3LG1*, encoding the ligand for NKp30, emerged as an important mediator of NK cell cytotoxicity against CML K562 and AML MOLM14 cells (Figure 4B-C) and was coherently highly expressed in NK-sensitive myeloid leukemias in the PRISM studies (Figure 3E). Other NK cell activating ligands that were depleted in tumor cells surviving NK cell treatment included *CD58* in K562 cells, *CD48* in ALL NALM6 and myeloma MM1.S cells, as well as *ULBP2* in NALM6 cells (Figure 4B-C). Activating receptor ligands *NCR3LG1* and *ULBP1* were concordantly among the genes most highly correlated with NK cell sensitivity in the PRISM studies (Figure 3E). Altogether, these findings suggest that while inhibitory signals by MHC-I and death receptor-mediated apoptosis are common across cell lines and cancer types, distinct activating receptor ligands promote NK cell cytotoxicity depending on the cell line.

Several adhesion molecules regulated the response to NK cell cytotoxicity. Beyond the established LFA-1 ligand *ICAM1*, which promoted NK cell killing of several cell lines, we also identified other adhesion molecules not previously associated with NK cell function, including *SPN* (*CD43*), the P-selectin ligand *SELPLG* (*PSGL1*), and *CD44*, which promoted resistance to NK cell killing. It has been reported that PSGL1, which was a hit in MM.1S, requires O-glycan fucosylation to become functional (Harjunpää et al., 2019). Interestingly, other fucosylation-related genes were identified regulating response to NK cell cytotoxicity in the LP1 cell line. The engagement of LP1 with NK cells was associated with a strong depletion of sgRNAs for genes essential for fucosylation including *FUT8, GMDS* and *SLC35C1* (Schneider et al., 2017), while the overexpression of terminal-fucosylation gene *FUT4* induced resistance to NK cell killing in the same cell line (Figure S3A). Overexpression of the mucin genes *MUC1* and *MUC21* conferred resistance to NK cells in MM lines. Altogether, these data point to a potential role of glycoproteins and surface ligand fucosylation for promoting resistance to NK cells.

A broad collection of transcriptional regulators and chromatin modifiers regulated NK cell response in hematologic malignancies. For example, loss of *ARID1A*, a member of the SWI/SNF chromatin remodeling complex, conferred resistance to NK cells in four cell lines (Figure 4B). In particular, the MM line MM1.S exhibited decreased response to NK cell treatment when *ARID1A* expression was lost, while it became significantly more sensitive when the same gene was overexpressed in the GOF screen. Several other regulators of gene expression also influenced sensitivity to NK cells, including the erythroid transcription factors *GFI1B* and *TAL1* in K562 CML cells, as well as the m6A RNA methylation modifier *YTHDF2* and the transcription factors *CMIP, FOXA1, PCGF5, RBBP4, IRF4, MYB*, and *MSI2* in other cell lines.

Other genes not previously linked to NK cell resistance included *SPPL3* in NALM6. LOF of the peptidase and positive regulator of HLA-I presentation *SPPL3* (Jongsma et al., 2020) conferred resistance also against CAR T cells, suggesting a mechanism related to general lymphocyte cytotoxicity instead of an NK cell-specific effect. In addition, our genome-scale screens identified the NF-κB negative regulator *NFKBIA*, the G-protein alpha subunit *GNA13*, and the antiviral gene *ZC3HAV1*, which was identified in both LOF and GOF screens in the MM cell line MM1.S.

We validated results from the genome-scale screens using a focused subgenome-scale sgRNA library as well as by targeting individual genes in select cell lines (Figures 4D-4E and S4A). These experiments largely reproduced the findings from the genome-scale screens, including the above-mentioned genes not previously linked to NK cell sensitivity, confirming their role in functionally regulating sensitivity to NK cell cytotoxicity.

### Genetic alterations and transcriptional regulation of NK cell susceptibility genes in cancer cells from patients with blood cancers

Alterations in CRISPR screen hit genes in cancer cells from patients with hematological malignancies could influence the efficacy of both the endogenous NK cell response and NK cell-based immunotherapies. To explore this, we first searched for somatic mutations in public datasets of AML, ALL, MM, DLBCL, and CLL. Somatic mutations occurred in the MHC-I complex subunit gene *B2M*, NF-κB signaling genes (*NFKBIA, BIRC3*), transcription factors and epigenetic modifiers (*ARID1A, IRF4, STAG2, MYB*), the CD2 ligand *CD58*, the extrinsic apoptosis mediator *CASP8*, as well as other genes such as *GNA13* (Figures 4F and S4B; Table S3). Many of these mutations were particularly prevalent in DLBCL (Figure 4F).

In addition to mutations, we investigated potential mechanisms regulating the expression of the CRISPR screen hits, including copy number alterations (CNAs) and DNA methylation, using multi-omics data from MM (CoMMpass), DLBCL (Reddy et al., 2017), and AML (TCGA, (2013)) (Figure S4C-D). Losses or deletions of *TRAF2* in MM and *JAK1, CD58*, and *ARID1A* in DLBCL were associated with reduced expression of the respective genes (Figure S4D; Table S3). CRISPR screen hits whose expression negatively correlated with DNA methylation of the transcription start site region included *TNFRSF1B, ULBP1*, and *ULBP3* in TCGA AML and *TNFRSF1B, ULBP1*, and *PVR* in TCGA DLBCL (Figure S4E; Table S3). AML patients with monocytic or myelomonocytic leukemia had the highest *TNFRSF1B* expression, indicating that both DNA methylation and cell type-specific transcriptional regulation can influence the expression of NK cell susceptibility genes such as *TNFRSF1B* (Figure S4E).

Regulators of NK cell sensitivity expressed exclusively or preferentially in blood cancers could represent blood cancer-specific NK cell regulators with potential for therapeutic targeting. Across all CCLE cell lines, we found the expression of *CD48, SPN, RHOH, MYB, SELPLG*, and *TNFRSF1B* as highly selective for blood cancers (Figures 4G and S4F). In solid tumor cell lines, these genes were highly methylated, indicating strong lineage-specific regulation of expression (Figure S4F). In contrast, the DNAM-1/CD226 and TIGIT ligand *PVR* and the NKG2D ligand *ULBP3* were enriched in solid tumors, although myeloid malignancies expressed *PVR* and T cell lymphomas (TCL) expressed *ULBP3* (Figures 4G and S4F). These expression patterns were also evident in primary patient samples in TCGA data (Figure S4F) as well as in normal healthy tissues (Figure S4G), indicating cell type, rather than oncogenic transformation, as the driver of the observed differences. Given their inhibitory function on NK cell cytotoxicity, genes such as *SELPLG, SPN*, and *MYB* therefore represent examples of blood-cancer-specific NK cell regulators that may present new therapeutic targets.

### Integration of CRISPR and PRISM screens reveals cancer subtype-specific NK cell evasion mechanisms

To find out if the mechanisms identified in our CRISPR screens could explain differential NK cell sensitivity of cell lines observed in the PRISM studies, we investigated whether transcript levels of genes (based on CCLE data) identified as hits in CRISPR screens correlated with the PRISM AUC. Across cancer types, high expression of genes including *NCR3LG1, ULBP1*, and *PVR* correlated with sensitivity, while *B2M, NLRC5, TAP1, CD44*, and *MSI2* correlated with resistance, indicating that these genes contribute to the differential NK cell sensitivity across blood cancer types (Figures 5A and S5A; Table S5). Stratified by cancer type, NK cell sensitivity correlated with high expression of *FAS, PVR* and *ULBP1* in T-ALL, *NCR3LG1* in B cell lymphoma (BCL), and *PVR* in B-ALL (Figure 5A), indicating heterogeneity in these activating ligands within certain cancer types in addition to their heterogeneity across cancers. The cancer type-specific analysis also identified resistance genes with potential preferential roles in individual neoplasms, including *NFKBIA* in BCL, *CFLAR* in MM, and *SPPL3* in B-ALL. Analysis of genetic signatures recurrently associated with NK cell resistance across all hematologic neoplasms revealed that the IFNγ response gene set retained its association with high PRISM AUC in AML, MM, and BCL, and a similar finding was observed for TNF alpha signaling via NF-κB in MM and BCL (Figure S5B). The core NK cell response signature and HLA I score correlated similarly with NK cell resistance only in BCL and MM (Figures S5C-S5D).

**Figure 5.**
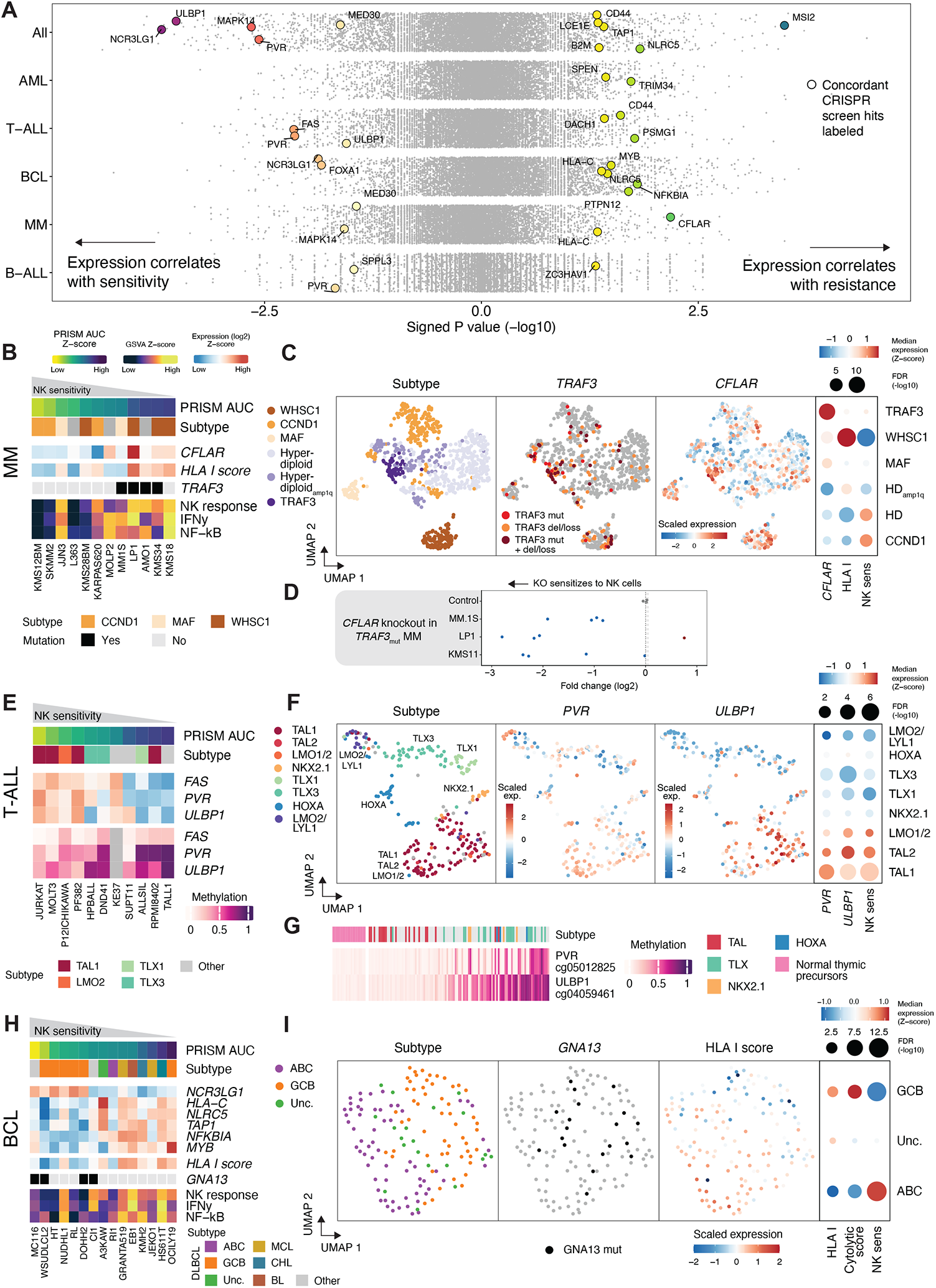
Integration of CRISPR and PRISM screens reveals cancer subtype-specific NK cell evasion mechanisms. (A) Gene expression correlations with sensitivity to NK cells (AUC) in PRISM shown as signed p values of the Spearman’s rank correlations using CCLE data. CRISPR screen hit genes (p < 0.0001) showing a correlation with PRISM AUC (p < 0.05) and with a correlation to the same direction as in CRISPR screens (e.g. lower expression correlates with NK cell sensitivity in PRISM and CRISPR-mediated silencing sensitizes to NK cells) are colored and labeled. Shown are correlations across all cell lines and within cell lines of individual cancer types. Dot color indicates the signed p value. (B) Heatmap of MM cell lines ordered by sensitivity to NK cells (PRISM AUC). Genetic subtypes, *CFLAR* expression, HLA I score, *TRAF3* mutation status, and GSVA scores of the core NK cell response (NK response), Hallmark interferon gamma response (IFNγ) and Hallmark TNFA signaling via NF-κB (NF-kB) gene sets are shown. (C) UMAPs of MM transcriptomic data from CoMMpass (n = 767). Genetic subtypes, *TRAF3* alterations, and *CFLAR* expression are colored on the plots. Dot plot on the right shows the median expression of *CFLAR*, HLA I score, and NK sensitivity signature (50 genes most significantly correlated with PRISM AUC in MM) across CoMMpass subtypes. Dot size indicates significance of differential expression between the indicated subtype and all other subtypes. (D) Effect of *CFLAR* knockout in genome-wide CRISPR screens in *TRAF3*-mutated MM cell lines. Log2 fold change between NK-treated and untreated cells is shown for each guide RNA. Olfactory receptor (OR) genes were used as a control and shown in gray is the average of all OR genes sgRNAs log2 fold change, while the dashed lines represent the 95% confidence interval. (E) Heatmap of T-ALL cell lines ordered by sensitivity to NK cells (PRISM AUC). Genetic subtypes, *FAS, PVR, ULBP1*, and *ULBP2* expression, and *FAS, PVR* and *ULBP1* methylation are shown. Color keys are shown above panel B. (F) UMAPs of T-ALL transcriptomic data from Liu et al. (n = 262). Genetic subtypes and *PVR* and *ULBP1* expressions are colored on the plots. Dot plot on the right shows the median expressions of *PVR, ULBP1*, and NK sensitivity signatures (50 genes most significantly correlated with PRISM AUC in T-ALL) across T-ALL subtypes. Dot size indicates significance of differential expression between the indicated subtype and all other subtypes. (G) Heatmap of *PVR* and *ULBP1* methylation in T-ALL patients (n = 109) and healthy controls (n = 20) from Roels et al. (GSE155333). The methylation probes with highest variance across the samples are shown. Samples are ordered based on average methylation of the two genes. Healthy thymocytes are shown on the left as comparison. Genetic subtypes are shown above according to Roels et al. (H) Heatmap of B cell lymphoma (BCL) cell lines ordered by sensitivity to NK cells (PRISM AUC). Lymphoma subtypes, expression of CRISPR hits correlating with PRISM AUC, HLA I score, *GNA13* mutations, and GSVA scores of the core NK cell response (NK response), Hallmark interferon gamma response (IFNγ) and Hallmark TNFA signaling via NF-κB (NF-kB) gene sets are shown. Color keys are shown above panel B. (I) UMAPs of DLBCL transcriptomic data from Chapuy et al. (n = 137). Cell-of-origin subtypes, *GNA13* mutations, and HLA I score are colored on the plots. Dot plot on the right shows the median HLA I score, cytolytic score, and NK sensitivity signature (50 genes most significantly correlated with PRISM AUC in BCL) across DLBCL subtypes. Dot size indicates significance of differential expression between the indicated subtype and all other subtypes. See also Figure S5 and Table S5.

We next asked whether genetic subtypes, mutations, or other factors could explain the heterogeneous expression of the NK cell susceptibility genes leading to variation in sensitivity within cancer types. In MM, among the genes most highly correlated with NK cell resistance were *CFLAR* encoding c-FLIP, a suppressor of death receptor-mediated apoptosis, and MHC-I genes (Figure 5B). MM cell lines harboring the t(4;14) (WHSC1) translocation tended to be more resistant, while the t(11;14) (CCND1) subtype tended to be more sensitive. Moreover, NK cell resistance was associated with inactivating mutations in *TRAF3*, known to induce activation of non-canonical NF-κB signaling (Keats et al., 2007; Liao et al., 2004). As *CFLAR* is a known NF-κB target gene (Micheau et al., 2001), *TRAF3* mutations could confer NK cell resistance by inducing *CFLAR*, resulting in impaired death receptor-mediated apoptosis.

We subsequently investigated whether these connections could also be found in patients using the CoMMpass data. We devised an NK cell sensitivity signature comprising genes whose expression correlated with sensitivity in the cell lines to identify patient subgroups with a molecular profile matching either NK-sensitive or NK-resistant cell lines. In agreement with the cell line data, the sensitivity signature was high in the CCND1 patient subgroup and low in the WHSC1 subgroup (Figure 5C), indicating that the cell lines faithfully represent molecular subtypes found in patients. *TRAF3* alterations, including nonsynonymous mutations, deletions, and losses, occurred both in a distinct *TRAF3*-altered cluster and in a subset of patients with *WHSC1* translocations, consistent with the cell lines where *TRAF3* mutations and *WHSC1* translocations often co-occured (Figure 5C). *CFLAR* expression was enriched in patients with *TRAF3* alterations belonging to both of these groups (Figures 5C and S5E). In addition, MHC-I expression was enriched in the WHSC1 subgroup, corroborating findings in the cell lines (Figures 5C and S5F). The functional role of *CFLAR* was validated in *TRAF3* mutated cell lines MM.1S, KMS11 and LP1, in which LOF of *CFLAR* induced increased response to NK cell attack (Figure 5D). These results indicate that *TRAF3* and *WHSC1* alterations confer an NK cell immune evasion phenotype in MM.

In T-ALL, NK cell sensitivity correlated with expression of the death receptor *FAS*, the DNAM-1 ligand *PVR* and the NKG2D ligand *ULBP1* (Figure 5E). The sensitive cell lines belonged to the TAL1 and LMO2 genetic subtypes representing late cortical differentiation (Ferrando et al., 2002) (Figure 5E). In contrast, other T-ALL lines showed low expression and high DNA methylation of *PVR* and *ULBP1*. Consistently, in patient genomic data from T-ALL (Liu et al., 2017), we found *PVR* and *ULBP1* expression enriched in the TAL1, TAL2, and LMO2 genetic subtypes corresponding to the late cortical stage (Figure 5F). Conversely, other subtypes including the TLX1 and TLX3 representing the early cortical stage and LMO2/LYL1 representing a more immature stage showed low expression of *FAS, PVR* and *ULBP1* (Figures 5F and S5G). *PVR* and *ULBP1* were unmethylated in patients with the TAL subtype and healthy T cells at all maturation stages, but showed increased methylation in other subtypes, suggesting that epigenetic control by DNA methylation may underlie the cancer-specific silencing of *PVR* and *ULBP1* in immature and early cortical T-ALL (Figure 5G) (Roels et al., 2020). Together, these findings suggest that the genetic background and differentiation stage of T-ALL can enable evasion from NK cells through decreased *PVR* and *ULBP1* expression in immature subtypes.

DLBCL cell lines of germinal center B-cell (GCB) cell-of-origin tended to be more sensitive to NK cells, whereas other BCL subtypes including MCL, CHL, Burkitt lymphoma (BL), and activated B cell (ABC) were less sensitive (Figure 5H). Expression of several CRISPR hits correlated with sensitivity, such as *NCR3LG1*, or with resistance, including *NLRC5, TAP1, NFKBIA, MYB, IRF4*, and MHC-I, also consistent with the IFNγ and NF-κB response gene sets being enriched in the resistant BCL cell lines. Mutations in *GNA13* correlated with NK cell sensitivity, in line with our CRISPR screen data showing sensitization to NK cells upon *GNA13* disruption (Figures 4C-4D). In DLBCL patients, the PRISM-derived NK cell sensitivity signature was consistently enriched in the GCB subtype, which harbored *GNA13* mutations and showed decreased expression of *NLRC5*, MHC-I genes, *TAP1, NFKBIA*, and *MYB* compared to ABC tumors (Figures 5I and S5H).

### Single-cell transcriptomics CRISPR screens reveal NK cell evasion mechanisms of genome-scale screen hits

We reasoned that the transcriptomic changes induced by perturbing genes identified in the CRISPR screens could reveal how these genes influence sensitivity to NK cells. We therefore performed pooled CRISPR screens with a single-cell transcriptome readout using the CROP-seq platform (Datlinger et al., 2017). We selected highly scoring hits from the genome-scale screens with a focus on likely transcriptional regulators, such as transcription factors and signaling molecules, as well as select NK cell ligands on tumor cells. We generated pools of cells expressing sgRNAs targeting hits in each cell line, including K562 (CML), SUDHL4 (DLBCL), NALM6 (B-ALL), MM1.S (MM), and LP1 (MM) and exposed the cells to NK cells at 1:16 or 1:4 effector-to-target ratios for 24 h or left untreated, followed by scRNA-seq and sgRNA assignment to cells (Figure 6A). In addition to the knockout CROP-seq screens, we performed a CRISPR activation (CRISPRa) CROP-seq analysis in MM1.S cells. From the six single-cell screens, we obtained a total of 118,968 cells with an assigned sgRNA, with three sgRNAs targeting each of the 65 perturbed genes and on average 128 cells representing each sgRNA (Table S5).

**Figure 6.**
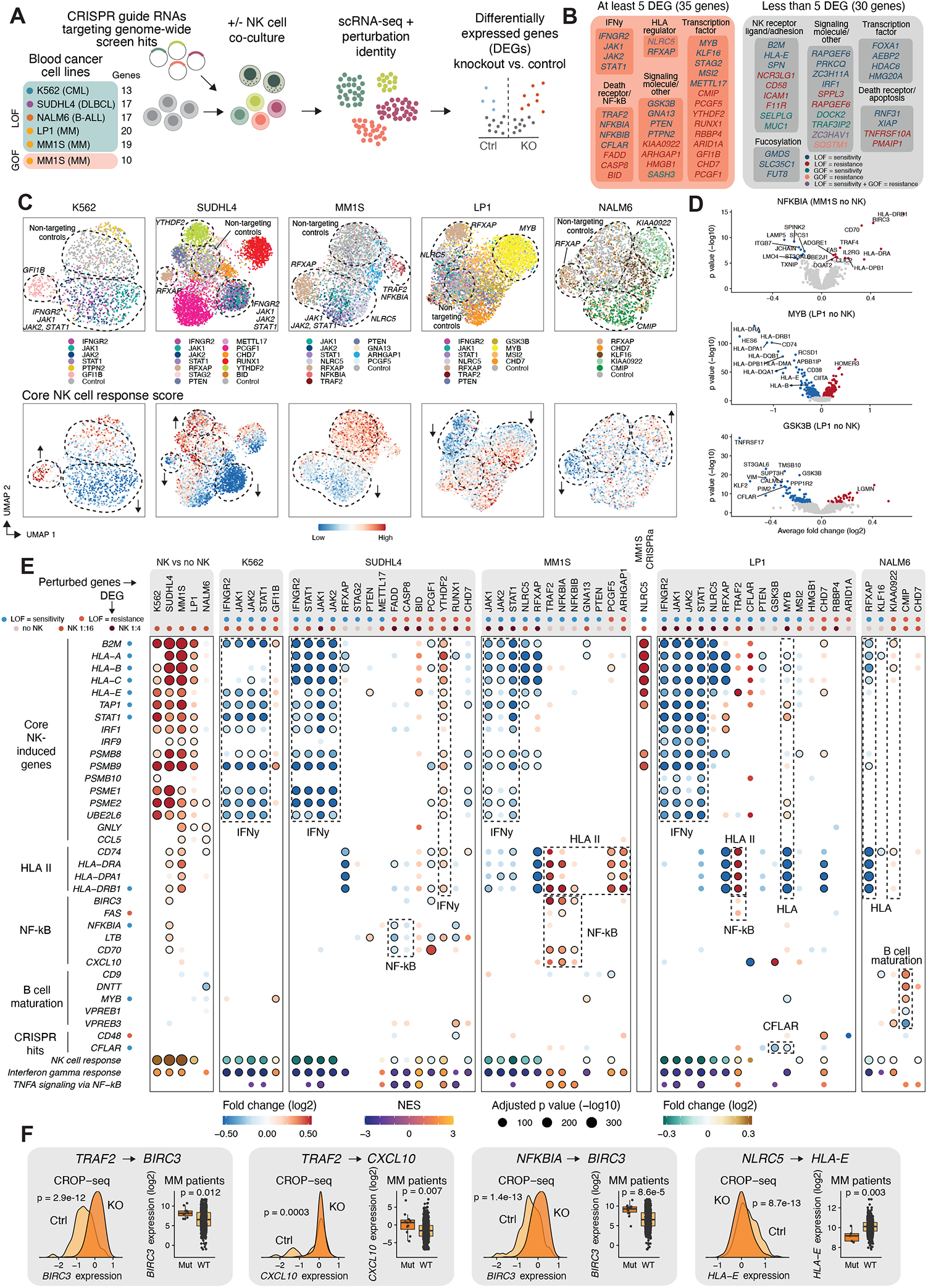
Single-cell transcriptomics CRISPR screens of cancer cell-intrinsic NK cell sensitivity regulators. (A) Single-cell CRISPR screening (CROP-seq) workflow. (B) Genes targeted in CROP-seq experiments, divided into groups with at least or less than 5 differentially expressed genes (DEGs) compared to non-targeting control cells in any of the conditions (no NK, NK 1:16, or NK 1:4). (C) UMAP visualizations of single-cell CRISPR screen data after running linear discriminant analysis in the indicated cell lines at 1:16 effector-to-target ratio. Cells classified as knockout or non-targeting (Control) by mixscape are shown. Perturbed genes (top row) and core NK cell response score (bottom row) are colored on the plots. (D) Volcano plots of differentially expressed genes with selected perturbations compared to control sgRNA-expressing cells. Red dots indicate genes with significantly (Bonferroni-adjusted p < 0.05) higher and blue dots lower expression in the CRISPR-targeted cells compared to control. (E) Dot plot of genes differentially expressed in cancer cells (rows) where the indicated genes (columns) are perturbed compared to cells expressing control sgRNAs. Perturbed genes with at least 5 DEGs are shown. For each perturbation, the condition (either NK 1:4, NK 1:16, or no NK) with the most significant differential expression is shown. Color indicates log2 fold change between conditions and dot size indicates the negative log10 adjusted p value. Only dots where p < 0.05 are shown, and circled dots indicate adjusted p value < 0.05. Enrichment or depletion of genes in genome-scale CRISPR screens are shown as colored dots both for the perturbations and perturbed genes. Effector:target (E:T) ratio for each perturbation is shown as colored dots above the plot. Selected molecular processes regulated by the perturbed genes are highlighted using dotted lines. (F) Examples of transcriptional changes induced by single-cell CRISPR screen perturbations with consistent changes observed in MM patients harboring mutations in the same genes in the CoMMpass data. Density plots of gene expression in scRNA-seq data in perturbed (dark yellow) and control cells (light yellow) are shown in the left column, and box plots of gene expression in patient RNA-seq data in mutated (‘Mut’) and non-mutated (‘WT’) samples are shown in the right column. Boxes indicate IQR with a line at the median. Whiskers represent the min and max values at most 1.5 IQR from the quartiles. See also Figures S6 and S7 and Table S6.

We analyzed differentially expressed genes between malignant cells harboring each perturbation and those carrying control sgRNAs, both with and without NK cell exposure. Out of the 65 perturbed genes, 30 showed no substantial transcriptomic changes, resulting in less than 5 differentially expressed genes in each perturbation (Figures 6B and S6A). These included genes encoding cell-surface proteins, such as *NCR3LG1, CD58, ICAM1, SPN, HLA-E*, or *MUC1*, suggesting that their physical interaction with NK cell surface molecules is the main mechanism mediating their effect on NK cell cytotoxicity, without other cancer cell-intrinsic molecular changes induced by the binding. For the 35 perturbations with a transcriptomic phenotype, we examined the common and distinct patterns by comparing the overlap of the differentially expressed genes (Figure S6B-C) and by using UMAP dimensionality reduction after performing linear discriminant analysis (Papalexi et al., 2021) (Figures 6C and S6D). Perturbations targeting IFNγ signaling mediators (*IFNGR2, JAK1, JAK2, STAT1*) grouped together in the UMAP reduced space, as did those targeting NF-κB regulators (*TRAF2, NFKBIA, NFKBIB*), consistent with common transcriptional changes induced by perturbing genes of the same pathway. In contrast, most other perturbations grouped individually, indicating distinct transcriptomic phenotypes. Some perturbations, including those targeting IFNγ and death receptor signaling mediators, induced substantial transcriptomic changes only in the presence of NK cells (Figures S6A and S6D). Several perturbations influenced the core NK cell response reflecting MHC-I genes and IFNγ signaling (Figures 6C and 6E), suggesting that these genes may regulate sensitivity to NK cells by influencing the transcriptomic response to NK cell attack.

We reasoned that if perturbing a CRISPR screen hit were to influence the expression of other hits, such observations could provide functional links between the former gene and the regulation of NK cell activity. Indeed, LOF of genes encoding IFNγ signaling mediators (*IFNGR2, JAK1, JAK2, STAT1*) in multiple cell lines prevented the NK-cell driven induction of the MHC-I complex genes, such as *B2M, HLA-E, HLA-A, HLA-B*, and *HLA-C*, as well as *TAP1* and *TAPBP* (Figures 6C, 6E, S6C, S6D, and S7A). The defective activation of *HLA-E* and other MHC-I genes therefore explains the sensitization to NK cells by disruption of the IFNγ-JAK-STAT components, consistent with an IFNγ-mediated negative feedback loop enabling target cell evasion from NK cells. Whereas LOF of the IFNγ signaling mediators showed a phenotype only in the presence of NK cells (Figures S6A and S7B), silencing of the HLA gene transactivator complex components *NLRC5* and *RFXAP* downregulated MHC-I genes both with and without NK cell exposure (Figure S7B).

Several genes which promoted NK cell killing, including *GFI1B* in K562, *YTHDF2* and *BID* in SUDHL4, *PCGF5* in MM1.S, and *KIAA0922* in NALM6, emerged as novel negative regulators of IFNγ signaling and MHC-I expression (Figures 6C, 6E, and S6B-S6D), providing a potential mechanism for the NK cell resistance conferred by their LOF. Conversely, LOF of *MYB* in LP1 cells reduced both MHC-I and MHC-II expression, indicating MYB as a positive regulator of antigen-presenting genes (Figures 6C-E, S6B, S6D, and S7B). MHC-I genes and IFNγ response were in general among the most recurrent differentially expressed genes and pathways across all perturbations, underlining the role of MHC-I and IFNγ regulators in controlling sensitivity to NK cells (Figure S6C).

To confirm some of the observed effects on MHC-I at the surface protein level and to discover additional ones not relying on transcriptional regulation, we integrated our CRISPR hits with MHC-I regulators identified by Dersh et al. (Dersh et al., 2021) (Figure S7D). Besides the known MHC-I components and *JAK1* concordant across the two sets of screens, *ARID1A* was identified as a negative regulator of MHC-I in the BJAB cell line and concordantly conferred resistance to NK cell attack in four of our genome-scale screens. In contrast, *TRAF2* and *CFLAR* had discordant effects in the two sets of screens, being negative regulators of MHC-I in Dersh cell lines and NK cell sensitizers in our MM screens, showing that this latter effect occurs through a different mechanism, overriding the potential HLA-I modulation.

Among potential mechanisms unrelated to MHC-I regulation, disruption of *NFKBIA* and *TRAF2* in MM1.S led to increased expression of the death receptor *FAS*, essential for susceptibility to NK cells in the MM1.S CRISPR screen (Figures 6D-6E and S7B). *GSK3B* and *MYB* LOF in LP1 downregulated *CFLAR*, a negative regulator of death receptor signaling and NK cell resistance gene identified in the CRISPR screens (Figures 6D-6E and S7B). *CMIP* disruption in NALM6 reduced the expression of *CD48*, ligand for the activating receptor 2B4 and essential for NK cell killing of NALM6 (Figures 6E and S7B). These regulatory networks between genes identified in the CRISPR screens provide mechanistic explanations for the observed resistance of sensitization to NK cell cytotoxicity unrelated to MHC-I regulation.

In addition to regulation of CRISPR screen hits, other perturbation-induced transcriptomic phenotypes could influence the interaction of the cancer cells with NK cells. In SUDHL4 cells, silencing of the death receptor apoptosis mediators *FADD* or *CASP8* inhibited the NK cell-induced NF-κB activation (Figures 6E and S7C). In addition to mediating apoptotic signals, FADD and CASP8 thus appear to regulate the transcriptomic response to NK cell attack (Henry and Martin, 2017; Kreuz et al., 2004). In contrast, silencing of *TRAF2, NFKBIA*, or *NFKBIB* in MM1.S cells induced NF-κB signaling (Figures 6D-E and S7A). This signature included known NF-κB targets such as *BIRC3, CD70*, and the chemokines *CXCL10* and *CCL5* (Annunziata et al., 2007; Herishanu et al., 2011), which may further increase immune reactivity through recruitment of T and NK cells (Loetscher et al., 1996a, 1996b; Schall et al., 1990; Taub et al., 1993). Silencing of *CMIP* (c-MAF interacting protein) in the pre-B-ALL cell line NALM6 led to increased expression of *DNTT* (*TdT*) and *CD9*, markers of early pro-B cells, and concomitant downregulation of genes expressed in more differentiated pro-B cells, including *CD79A, VPREB1*, and *VPREB3* (Figures 6E, S6D, and S7A). A more immature B cell state controlled by *CMIP* therefore may drive resistance to NK cell killing. Overall, the single-cell CRISPR perturbation data offer a resource of the phenotypic consequences of altering genes regulating NK cell-cancer cell interactions, providing mechanistic explanations and testable hypotheses for further exploration.

Given that NK cell susceptibility genes were mutated in various hematological malignancies, we asked whether these mutations would result in similar transcriptomic alterations in patient cells as observed in the single-cell CRISPR screens. We compared the differentially expressed genes of each scRNA-seq perturbation with the differentially expressed genes between patients with and without mutations in the same gene using multi-omic data from MM (CoMMpass), DLBCL (Reddy et al., 2017), and AML (TCGA) (Figures 6F and S7E). MM patients with mutations in the NF-κB negative regulators *TRAF2* and *NFKBIA* expressed higher levels of NF-κB target genes also identified experimentally by CROP-seq (Figures 6F and S7E). Moreover, MM patients with *NLRC5* mutations had lower *HLA-E* expression consistent with CROP-seq data (Figure 6F, S7E), indicating that although rare, *NLRC5* mutations in MM may result in an NK-sensitive phenotype (Figure 6E). These findings provide evidence that several regulatory mechanisms identified by the single-cell CRISPR screens operate in patient cells and therefore can influence susceptibility to NK cells *in vivo*.

### Cancer-cell intrinsic perturbations modulate NK cell transcriptomic states

Finally, we asked if perturbing CRISPR screen hits in the cancer cells could influence the NK cell activation states we originally observed upon co-culture with various cancer cell lines (Figure 1C). We co-cultured *ex vivo*-expanded NK cells with three cell lines (K562, SUDHL4, and NALM6) expressing individual sgRNAs targeting 14 different genes (Figure 7A). After 24 h co-culture at 1:1 effector-to-target ratio, leading to elimination of most target cells, we performed multiplexed scRNA-seq on the NK cells using cell hashing. To quantify the effects of the target cell knockouts on NK cell activation, we computed an activation score for each NK cell, comprising 50 genes most significantly enriched in the activated NK cell cluster (cluster 2). Comparison of the activation scores of NK cells co-cultured with cancer cells harboring different perturbations to those cultured with non-targeting control sgRNA-expressing cells revealed that *CD58* LOF in K562 strongly reduced NK cell activation (Figure 7B). In addition to *CD58*, other resistance-inducing perturbations including *CMIP* and *SPPL3* LOF in NALM6 had significant inhibitory effects. In contrast, *JAK1* LOF induced stronger NK cell activation, consistent with the sensitizing effect (Figure 7B). Interestingly, *KCNH2* LOF induced resistance to NK cell cytotoxicity but still showed an activating effect on NK cells.

**Figure 7.**
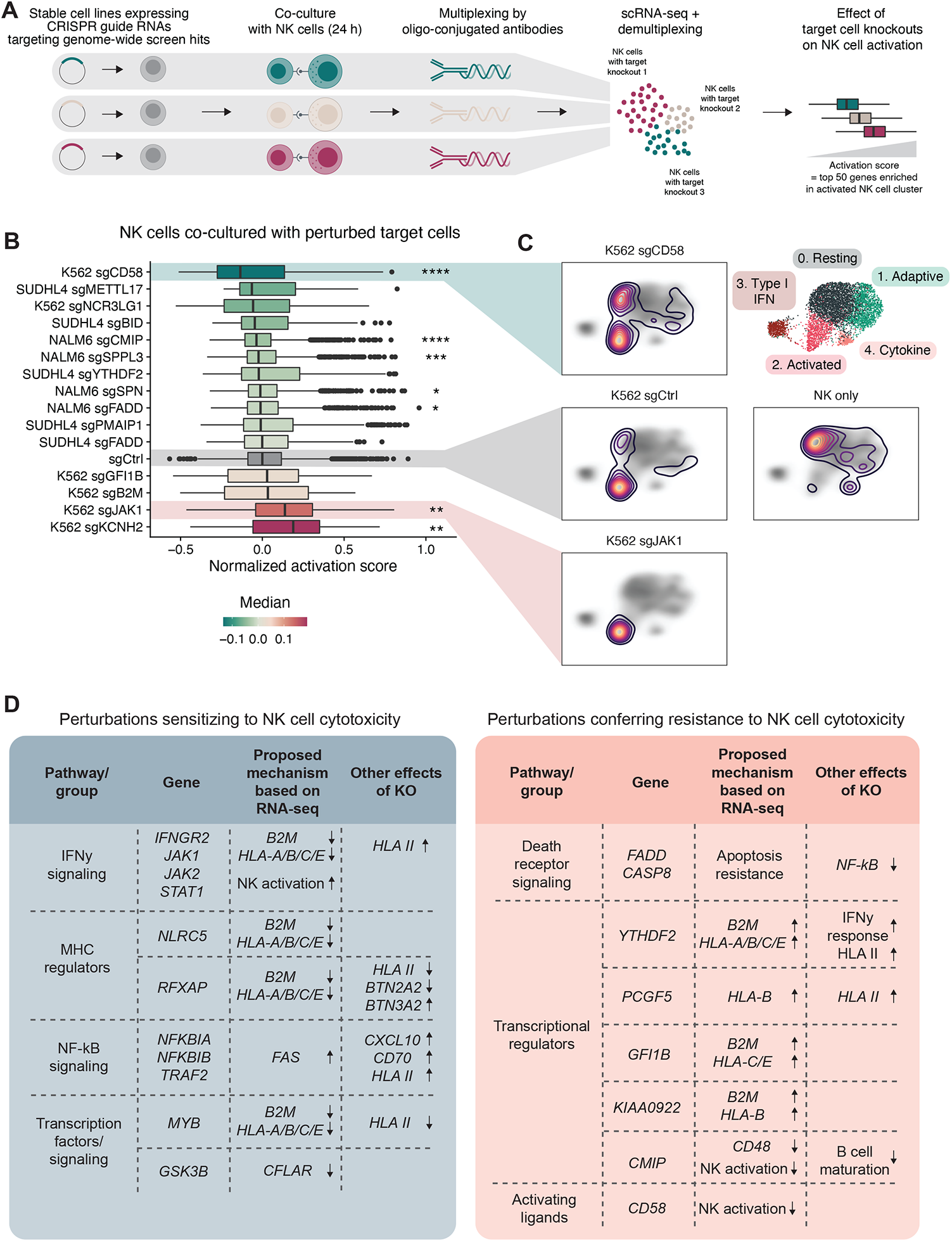
Effects of cancer cell perturbations on NK cell activation states. (A) Workflow of identification of NK cell responses to cancer cells carrying different perturbations using single-cell transcriptomics. (B) Box plot of NK cell activation scores across different target cell perturbations. Activation scores comprising top 50 genes enriched in the activated NK cell cluster were normalized by subtracting the median activation score of the control sgRNA-expressing target cells of each cell line. P values between each perturbation and the cell line-specific control are obtained using Wilcoxon rank sum test with Benjamini-Hochberg adjustment. Only p values for significant pairs are shown (* < 0.05, ** < 0.01, *** < 0.001, **** < 0.0001). Boxes indicate IQR with a line at the median. Whiskers represent the min and max values at most 1.5 IQR from the quartiles. (C) UMAP visualizations of NK cells co-cultured with K562 cells expressing the indicated sgRNAs or NK cells cultured alone. The contour lines and their color indicate the density of NK cells in different regions of the UMAP reduced space. The gray shading in the background shows the density of all NK cells from Figure 1 and the clusters from Figure 1 are shown on the right as a reference. (D) Summary tables of findings from the single-cell transcriptomics assays on perturbation effects both in the target cells (CROP-seq) and in NK cells, including the proposed mechanism of induced sensitivity or resistance to NK cells and other transcriptional effects observed in knockout (KO) cells.

To visualize differences in NK cell states relative to those identified in the co-culture experiments using the 26 different cell lines, we mapped the NK cells from the present experiment to the UMAP dimensionality reduction from the previous experiment (Figure 7C). While most NK cells exposed to control K562 cells moved from the resting to the activated cluster, *CD58* LOF in K562 cells caused a substantial fraction to remain in the resting state. Conversely, *JAK1* LOF induced almost all NK cells to change to the activated state. These findings indicate a key role for the CD2-CD58 interaction in promoting the activated state and conversely for JAK1-mediated signaling in inhibiting the activation. The perturbation-induced transcriptional changes both in NK cells and in target cells thus offer plausible mechanisms for how the genes identified in CRISPR screens control sensitivity to NK cells and provide a comprehensive resource of other immunoregulatory effects (Figure 7D).

All the data from multiplex scRNA-seq co-culture experiments, genome-wide and single cell CRISPR screens and PRISM screens together with the molecular correlates are available for interactive exploration at https://immunogenomics.shinyapps.io/nkheme/.

## DISCUSSION

In this study, we systematically mapped the landscape of the interaction of NK cells with diverse types of tumor cells across hematological malignancies. We studied the phenotypic changes induced by the interaction of effector and target cells at single-cell resolution, profiled the sensitivity of different cancers and molecular subtypes to NK cells by PRISM profiling, and identified cancer cell genes and pathways influencing sensitivity to NK cells using CRISPR screens. Besides certain common core mechanisms across different diseases, a key finding emerging from the integration of these data is the heterogeneity of mechanisms influencing NK cell susceptibility between individual cancers driven by lineage and molecular subtypes of cancer. Our findings indicate a need to consider the cancer subtype and genetics for optimal tailoring of NK cell-based therapies for blood cancer patients.

The diverse adaptive, interaction-induced responses detected in our single-cell studies reflected the heterogeneity, as different cell lines induced in NK cells a transition towards distinct activated states that ranged from a full shift to very little or no changes at all. The NK cell clusters enriched upon tumor cell exposure emerged both in expanded NK cells and in unexpanded PBMC-derived NK cells, and included an activated state (cluster 2), a cluster with high type I IFN signature (cluster 3) possibly resembling the previously identified type I IFN-responding cells (Smith et al., 2020) and inflamed NK cells (Yang et al., 2019), and a cytokine-producing phenotype (cluster 4). The activated state included genes encoding receptors such as 4-1BB, OX-40, and CRTAM, shown also previously to be induced by NK cell activation (Baessler et al., 2010; Costanzo et al., 2018; Turaj et al., 2018). As the activated states correlated with increased cytotoxicity against target cells, interventions promoting the activated state such as agonistic antibodies for activating immune checkpoint receptors could improve NK cell immunotherapies. Some of the receptors induced upon transition to the activated state inhibit NK cell function, including TIM-3, TIGIT, and possibly also 4-1BB (Baessler et al., 2010) and GITR (Baltz et al., 2007). Blocking these inhibitory signals could further augment the function of NK cells recognizing their targets.

The gene signature recurrently induced in cancer cells in response to NK cell exposure reflecting IFNγ signaling and MHC-I is consistent with a negative feedback loop suggested in early studies of NK cell effects on target cells (Piontek et al., 1985; Trinchieri and Santoli, 1978). Our single-cell CRISPR data confirm the role of interferon signaling through JAK-STAT in driving the observed responses. The same transcriptomic signatures that were induced in cancer cells upon NK cell attack correlated with resistance to NK cells across blood cancer cell lines. Pre-existing activation of adaptive resistance pathways may therefore explain primary resistance of cancer cells to NK cells. Conversely, as defects in interferon signaling and antigen presentation cause resistance to T cell immunotherapies (Zaretsky et al., 2016) as suggested also by previous CRISPR studies, (Freeman et al., 2019; Sheffer et al., 2021) NK cells could offer an effective alternative; and a concrete opportunity for individualized use of NK cell therapies in patients whose blood cancer cells harbor genomic defects in this molecular cascade.

Our findings challenge the notion that expression of activating ligands for NK cells is a general feature of transformed cells (“altered-self”). Instead, the cancer type, lineage, and genomics appear to jointly define the expression patterns. For example, previous CRISPR screen studies of NK cell resistance in blood cancer cells performed in K562 cells identified *NCR3LG1* as essential for NK cell cytotoxicity (Pech et al., 2019; Zhuang et al., 2019). Our data confirm this finding but also provide key new insights that *NCR3LG1* appears important for effective NK cell killing particularly in myeloid leukemias, unlike several other types of hematological malignancies.

The variation in expression of activating ligands, including *NCR3LG1, PVR*, and *ULBP1*, translated into differences in sensitivity to NK cells, with AML being sensitive compared to pre-B-ALL as also previously suggested (Pende et al., 2005). The expression of activating receptor ligands and resulting sensitivity to NK cells may be particularly pronounced in more differentiated myeloid cells represented by most AML cell lines. Instead, a less differentiated phenotype can enable evasion from NK cells (Nowbakht et al., 2005; Paczulla et al., 2019). The lineage-dependent expression of genes encoding activating ligands such as *PVR* or apoptotic mediators such as *TNFRSF1B* both in myeloid malignancies and their normal counterparts implies that this expression pattern originates from normal hematopoietic differentiation rather than being a feature acquired upon transformation. Moreover, identification of previously unknown blood cancer-specific NK cell immune regulators including *SPN* and *SELPLG* and the lineage-driven expression of genes encoding known ligands such as *PVR* and *ULBP3* support that blood cancers may be configured to interact with NK cells in a way distinct from solid tumor cells, possibly because they originate from various immune cell types that naturally frequently interact with NK cells. Our CRISPR screens implicated many previously underappreciated gene classes in the regulation of NK cell responses, including protein fucosylation, mucins, and a range of transcriptional regulators, with potential relevance across blood cancers and solid tumors. As many of these were uncovered only in a subset of the cell lines, our data thus highlight the importance of performing large unbiased screens in diverse cancer types.

In addition to cancer types and lineages, our findings link several previously established molecular subtypes of blood cancers with characteristic genetic alterations to NK cell evasion mechanisms. The transcriptomic cluster in MM characterized by *TRAF3* alterations and activation of *CFLAR* and other NF-κB targets likely reflects the NF-κB cluster identified in early transcriptomic studies of MM (Broyl et al., 2010). Given that *CFLAR* appears to influence sensitivity of cancer cells not only to NK cells as indicated by our data but also T cells (Singh et al., 2020; Vredevoogd et al., 2019), this finding may have relevance to immunotherapy sensitivity beyond NK cells. The methylation of *PVR* and *ULBP1* in immature subtypes of T-ALL may be linked to the CpG island methylation phenotype (CIMP) found in non-TAL1-driven T-ALL (Borssén et al., 2013; Kimura et al., 2020; Roels et al., 2020), suggesting that one of the functions of CIMP in T-ALL may be enabling immune evasion from NK cells. Together, these findings begin to unravel the connections between cancer genomic subtypes and responsiveness to NK cells that have thus far remained largely unexplored.

While NK cell-mediated cytotoxicity depended on intact death receptor signaling in several cell lines, this pathway was not relevant in some cell lines, such as the classical NK cell target K562. Thus, heterogeneity appears to exist also in the apoptotic mechanisms even between NK-sensitive cell lines. The reliance on distinct apoptotic pathways in different cancers may influence the therapeutic approaches to sensitize tumors to NK cells, such as the recently proposed BH3 mimetics (Pan et al., 2022). Given that the death receptor pathway can also mediate bystander killing (Upadhyay et al., 2020), the differential sensitivity of different cancers to bystander killing may influence the efficacy of both NK and T cell immunotherapies.

Our study has several limitations. Although our studies included over 60 blood cancer cell lines, more rare types of blood cancers were represented by few or no samples. Moreover, our approach to investigate the mechanisms by which genes identified in CRISPR screens influence NK cell responsiveness using CROP-seq likely misses several mechanisms that do not operate primarily through altering gene expression. Furthermore, as many of the discovered mechanisms are dynamic, validating the results *in vivo* in patients would require multiple sampling during adoptive cell transfer to capture the changes occurring immediately when NK cells come in contact with the cancer cells.

In summary, our study provides a comprehensive picture of both the adaptive molecular changes in interacting NK cells and tumor cells as well as genetic mechanisms of response and resistance of blood cancer cells to NK cell cytotoxicity. These molecular profiles offer a resource that can inform efforts to develop NK cell immunotherapy strategies particularly in hematological malignancies, which are emerging as the primary clinical setting for the application of NK cell-based therapies.

## METHODS

### Cell lines

PL21, GDM1, and SKM1 were cultured in RPMI-1640 with 10% heat-inactivated fetal bovine serum (FBS), 2 mM L-glutamine, and 100 U/mL penicillin with 100 mg/mL streptomycin (PS). OCIM1 were cultured in IMDM (Gibco) with 10% FBS, 2 mM L-glutamine, and PS. All other cell lines were grown in RPMI-1640 with 10% FBS, 2 mM L-glutamine, and PS. All cultures were incubated at 37°C with 5% CO_2_.

PRISM cell line pools were cultured in phenol red-free RPMI 1640 with 20% FBS and PS.

KHYG1 were cultured in RPMI-1640 with 10% FBS (20% for first passage upon thawing as per manufacturer’s instructions), 1% PS and 100 IU/ml of human recombinant IL-2 (R&D Systems, 202-IL-050). Cells were used at low passage numbers, in order to avoid the outgrowth of growth factor independent subclones.

To generate Cas9-expressing K562, SUDHL4, NALM6, and MOLM14 cells, the cells were transduced with virus produced using the lentiCas9-EGFP plasmid (a gift from Phil Sharp & Feng Zhang, Addgene plasmid # 63592), single-cell sorted using a Sony SH800 cell sorter, and a clone with high and uniform EGFP expression was selected for screening. The MM.1S-Cas9+ cells were generated and kindly gifted by the laboratory of Dr Benjamin Ebert (DFCI). KMS11-Cas9+ cells and LP1-Cas9+ cells (transduced with pLX 311-Cas9 construct, Addgene plasmid # 96924) were obtained from the Broad Institute, as well as MM.1S-dCas9VP64, KMS11-dCas9/VP64 and LP1-dCas9/VP64 (transduced with lenti dCAS-VP64_Blast, Addgene plasmid # 61425).

Luciferase-expressing K562 cells were generated using the pLenti PGK V5-LUC Neo (w623-2) construct as previously described for NALM6 (Dufva et al., 2020a). The generation of luciferase-expressing SUDHL4 cells has been previously described (Dufva et al., 2020a).

All cell lines were STR profiled and tested for Mycoplasma using the MycoAlert kit (Lonza).

### Primary NK cell isolation and expansion

#### Expansion with feeder cells

NK cells were expanded using K562-mbIL21-41BBL feeder cells as previously described (Denman et al., 2012). Briefly, PBMCs were isolated from buffy coats of healthy donors using Ficoll-Paque gradient centrifugation. Five million PBMC were suspended in 40 ml R10 supplemented with 10 ng/ml recombinant human IL-2 (R&D Systems, 202-IL-050) together with 10 million K562-mbIL21-41BBL feeder cells irradiated with 100 Gy. Cells were passaged twice a week and feeder cells were added in a 1:1 ratio after 7 days. After 14 days of culture, NK cells were purified using the NK Cell Isolation Kit (Miltenyi) and frozen. NK cells from various donors were thawed and cultured for 5 days in R10 + IL-2 prior to genome-scale CRISPR screens (K562, MOLM14, SUDHL4, NALM6). For all multiplexed scRNA-seq and CROP-seq experiments, NK cells from the same donor were used and thawed and cultured 3 days prior to the experiments.

#### Expansion without feeder cells

PBMCs were isolated from consenting healthy donors. CD3+ cells were depleted using the negative selection cocktail RosetteSep™ (STEMCELL Technologies Inc.) according to the manufacturer’s instructions. CD3-negative PBMCs were then seeded in 6-well plates at a density of 1×10^6^ cells/ml in GMP SCGM media (CellGenix) with 10% heat-inactivated FBS, 1% Glutamax, 200 IU/ml of recombinant human IL-2 (R&D Systems, 202-IL-050) and expanded for 10 – 14 days. Expanded NK cells from PBMCs from the same donor (donor #9) were used in the genome-scale and focused-library screens with MM1S, LP1, and KMS11 cells and the PRISM screen. Flow cytometry was performed to verify primary NK cell viability, purity (anti-CD56-PECy7 and anti-CD3-FITC), and expression of p46 receptor (anti-NKp46-APC), surrogate marker of NK cell activity.

### Co-culture assays with multiplexed scRNA-seq readout

#### Experiments and scRNA-seq library preparation

For the experiment involving 26 different cell lines, cancer cells were plated at 500,000 cells/well on a 24-well plate and day 17 feeder cell-expanded NK cells (1:4 effector-to-target ratio) or NK cells directly extracted from PBMC from the same donor (1:2 effector-to-target ratio) or only R10 culture medium (targets only) were added, resulting in a total volume of 1 ml R10. Experiments were performed in two batches of 13 cell lines, and wells with only NK cells were included.

For the experiment involving CRISPR-targeted cell lines, day 17 feeder cell-expanded NK cells were used at an 1:1 effector-to-target ratio.

After 24 h in 37°C and 5% CO_2_, cells from each well were washed 2-3 times with 10 ml PBS, resuspended in 100 µl Cell Staining Buffer (BioLegend), 10 µl TruStain FcX blocking reagent (BioLegend) was added, and cells were blocked for 10 min. A unique TotalSeq-A hashing antibody (BioLegend) was added to each sample (1-2 µl/1-2 µg per sample) and cells were incubated for 30 min at +4°C covered from light. Cells were then washed 3-5 times with 3 ml staining buffer and samples were combined in 1 ml staining buffer, centrifuged, resuspended to PBS + 0.04% bovine serum albumin (BSA) and proceeded to scRNA-seq. The Chromium Single Cell 3’RNAseq run and library preparations were done using the 10x Genomics Chromium Next GEM Single Cell 3’ Gene Expression version 3.1 Dual Index chemistry with the modifications described in Stoeckius et al. (Stoeckius et al., 2018), https://cite-seq.com/ and according to the slightly improved protocol described in www.biolegend.com/en-us/protocols/totalseq-a-antibodies-and-cell-hashing-with-10x-single-cell-3-reagent-kit-v3-3-1-protocol. The 3’ GEX and Cell Hashing (multiplexing) libraries were sequenced using Illumina NovaSeq 6000 system using read lengths: 28bp (Read 1), 10bp (i7 Index), 10bp (i5 Index) and 90bp (Read 2).

#### Data analysis

Data preprocessing was performed using 10x Genomics Cell Ranger v6.0.2 pipelines. The ‘cellranger mkfastq’ was used to produce FASTQ files and ‘cellranger count’ to perform alignment, filtering, and UMI counting. The Illumina bcl2fastq v2.2.0 was used to run mkfastq and alignment was done against human genome GRCh38. Count matrix for hashtag oligonucleotides (HTO) was generated using the CITE-seq-Count-tool (DOI 10.5281/zenodo.2590196) (Stoeckius et al., 2018).

The R package Seurat (v4.0.4) (Stuart et al., 2019) was used for further scRNA-seq data processing. Cells with > 15% mitochondrial gene counts, > 50% or < 5% ribosomal gene transcripts, < 700 UMI counts, or < 300 or > 10,000 detected genes were filtered out. Hashtag oligonucleotide (HTO) demultiplexing to classify cells to samples was performed on centered log-ratio-normalized HTO UMI counts using the HTODemux function in Seurat with a positive quantile of 0.99. Sample IDs based on HTO data were transferred to transcriptome data and only cells classified as singlets based on HTODemux were considered for further analyses. After log-normalization, the highly variable genes were calculated with the FindVariableFeatures function using the “mean.var.plot” selection method in Seurat. Data were scaled and the effect of the cell cycle was corrected using the ScaleData function with scores assigned to each cell using the CellCycleScoring function with G2/M and S phase markers provided in Seurat. Clusters were defined using the FindClusters function with resolution set to 0.8 and cell types were annotated using SingleR (Aran et al., 2019). Clusters comprising NK cells were identified and subsequent analyses were focused only on expanded or PBMC NK cells (NK cell clusters) or cancer cells (all other clusters). The UMAP dimensionality reduction (McInnes et al., 2020) with default parameters was calculated using RunUMAP with the top 20 principal components (PCs).

For the analysis focusing on NK cells (either expanded or PBMC NK cells), data were re-scaled and the cell cycle effect and batch (resulting from performing the experiments in two batches) were corrected for using the ScaleData function in Seurat. Clusters were defined using the FindClusters function with resolution set to 0.3, and the UMAP was calculated from the top 20 PCs. Differentially expressed genes between clusters were obtained with a Student’s *t*-test followed by Bonferroni correction using the FindAllMarkers function in Seurat. Pseudotime analysis was performed using Slingshot (v2.2.0) (Street et al., 2018) on the precalculated UMAP coordinates, with cluster 0 (“Resting”) assigned as the start and cluster 2 (“Activated”) as the end.

For the analysis focusing on cancer cells, differentially expressed genes between NK cell-treated and untreated cells were obtained with a Student’s *t*-test followed by Bonferroni correction using the FindMarkers function in Seurat. Multiple testing correction was performed separately for each cell line comparison. For the differential expression analysis across all cell lines (Figure 2A), 1,000 cells were subsampled from the treated and untreated cells. For UMAP visualizations of each individual cell line, data were scaled, the cell cycle effect was regressed out using ScaleData, and small clusters comprising less than 5% of cells (representing misclassified other cell lines) were removed. The UMAP dimensionality reduction with default parameters was calculated from the top 20 principal components. The core NK cell response score was calculated using the AddModuleScore function in Seurat based on the 16 genes induced by co-culture with expanded NK cells in > 75% of the cell lines (*B2M, HLA-A, HLA-B, HLA-C, HLA-E, TAP1, STAT1, IRF1, IRF9, PSMB8, PSMB9, PSMB10, PSME1, PSME2, UBE2L6, GNLY*, and *CCL5*).

Ligand-receptor interactions were calculated using CellPhoneDB (Efremova et al., 2020) with default parameters from each cell type subsampled to the same number of cells. Interactions were calculated between all NK cell clusters and each cell line (both untreated and NK cell-treated), and the significant interactions (p < 0.05 permutation testing) based on CellPhoneDB were considered for downstream analyses.

To compute the activation scores for the NK cells co-cultured with CRISPR-targeted cell lines, the AddModuleScore function in Seurat was used on the 50 genes most significantly enriched in the activated cluster (cluster 2) in the cell line panel experiment. When NK cells cultured with target cells expressing two different sgRNAs targeting the same gene were available, the NK cells were pooled together for the analysis. Wilcoxon rank sum test was used to compare NK cells cultured with a gene-targeted cell line to those cultured with the same cell line expressing non-targeting control sgRNAs. The normalized activation scores were obtained by subtracting the median activation score of the same cell line control from the activation scores of the NK cells co-cultured with gene-targeted cell lines. The NK cells co-cultured with CRISPR-targeted cell lines were projected onto the previously computed UMAP visualization from the 26 cell line panel experiment using the FindTransferAnchors and MapQuery functions in Seurat.

### Pooled PRISM screen of NK cell cytotoxicity against DNA-barcoded cancer cell lines

PRISM is a platform that allows pooled screening of mixtures of cancer cell lines by labeling each cell line with 24-nucleotide barcodes as previously described (Yu et al., 2016). Briefly, 70 suspension blood cancer cell lines (Table S2) stably expressing DNA barcode sequences were seeded in 6-well plates in 8 experimental replicates per condition. The cells were incubated in 5 ml PRISM growth medium (RPMI-1640 without phenol red + 20% FBS + PS) for 24 h. At that point, primary NK cells were washed, resuspended in PRISM growth medium and added to the PRISM cells in 4 different E:T ratios 5:1, 2.5:1, 1.25:1, and 0.625:1 (1 ml/well). Control wells were added with the same volume of media only.

After 24 h co-culture, cells from each well were washed with PBS and incubated for 1 hour at 60°C in lysis buffer (1 ml per well), prepared using double-distilled water with 10% PCR buffer (20mM Tris-HCL PH 8.4, 50mM KCL), 0.45% NP40, 0.45% TWEEN and 10% proteinase K. Genomic DNA from cell lysate was amplified, PCR product was hybridized to Luminex beads with covalently attached antisense barcodes, and streptavidin-phycoerythrin addition, washing, and detection on Luminex FlexMap machines was performed as previously described (Yu et al., 2016).

Means of the eight experimental replicates of each cell line were calculated for each E:T ratio and percent viability values were obtained by dividing the mean of each E:T ratio with the mean of the untreated control for each cell line multiplied by 100. Area under the curve (AUC) values were calculated with the percent viability values using the AUC function in the DescTools (v0.99.43) R package. Three non-hematological cell lines included in the pool were removed from the analysis: gastric adenocarcinoma cell lines HUG1N and SNU1; Ewing sarcoma cell line CHLA57. Cell lines with incomplete data at all E:T ratios were similarly removed from the analysis, resulting in 63 cell lines.

### Genome-scale CRISPR/Cas9-based gene editing or gene activation screens

#### Production of viral particles

Brunello/Calabrese screens: Lenti-X-293T cells (Takara Bio) were plated in T-175 culture flasks (0.6×10^6^ cells/ml) in DMEM (Life Technologies) with 10% FBS for 24 h. After decanting the cell medium, OPTI-MEM (6 ml) and Lipofectamine 2000 (100 μl; Life Technologies) were added to each flask plus packaging plasmids psPAX2 (20 μg) and MD2.G (10 μg) and plasmid preps of the Brunello sgRNA library or Calabrese sgRNA library (20 μg per prep; lentiGuide-Puro). Plasmid preps for the Brunello and the Calabrese sgRNA libraries were purchased from Addgene (#73178 and #1000000111). The transfected Lenti-X-293T cells were incubated at 37ºC (20 min), topped up with fresh media (25 ml), and then refreshed again after 16 hours. Viral supernatants were collected after 24 h and stored at −80ºC prior to use.

GeCKOv2 screens: The genome-scale GeCKO v2 sgRNA library in the lentiGuide-Puro plasmid (Sanjana et al., 2014; Shalem et al., 2014) (a gift from Feng Zhang, Addgene # 1000000049) was amplified using Endura competent cells (Lucigen) according to instructions provided by the Zhang lab and Lucigen as previously described (Dufva et al., 2020a). To produce lentivirus, 10 µg of both A and B library plasmids were transfected into 293FT cells seeded on the previous day at 11.4 million cells/T-225 flask, together with 15 µg of psPAX2 and 10 µg of pCMV-VSV-G using 100 µl Lipofectamine 2000 (Thermo Fisher Scientific) and 200 µl of Plus Reagent (Thermo Fisher Scientific). After 6 h incubation, the culture medium was replaced with 30 ml of D10 containing 1% BSA. After 60 h, the viral supernatant was harvested, filtered using a 0.45 µm filter, and stored in −70°C.

#### Lentiviral transductions with sgRNA libraries

Brunello LOF screens: Tumor cell transductions were performed in batches of 5×10^7^ cells per library for three replicates. Cells were incubated (18 h) in cell medium containing polybrene (5 μg/ml; Santa Cruz Biotechnology), 10 mM HEPES (pH 7.4) (Gibco) and viral prep (30 ml) diluted 1:1. Transduced cells were cultured at an initial density of 1×10^6^ cells/ml and were treated with puromycin (1 μg/ml) for up to 5 days additional two days from transduction. After stable transduction, pooled cells were plated at 40×10^6^ cells per flask (T-175, 100 ml) to enable coverage of 500X and were sub-cultured at three- to four-day intervals to prevent confluence.

Calabrese GOF screens: Tumor cells were transduced in batches of 3×10^7^ cells per sub-library in triplicates. Cells were incubated (18h) in cell medium containing polybrene (4 μg/ml; Santa Cruz Biotechnology), 10mM HEPES (pH 7.4) (Gibco) and viral prep (30 ml) diluted 1:1. Transduced cells were cultured at an initial density of 1×10^6^ cells/mL and were treated with puromycin (1μg/mL) for up to 7 days additional two days from transduction. After stable transduction, pooled cells were plated at 30×10^6^ cells per flask (T-175, 100 ml) to enable coverage of 500X and were treated with primary NK cells in either duplicates or triplicates. The E:T ratio was selected to kill at least 50% of the tumor cells, according to a dose-response curve.

GeCKO v2 LOF screens: The amount of lentivirus used to transduce the cells was first optimized by transducing cells with a range of virus concentrations on a 12-well plate, where in each well 3 million cells were suspended in a total volume of 1 ml containing 0-1000 µl of GeCKO v2 library virus and 8 µg/ml Polybrene. The plate was centrifuged at room temperature at 800 g for 2 h after which virus 8 was washed away. The cells were treated with or without 0.5 µg/ml puromycin (Thermo Fisher Scientific) for 6 days starting 48 h post-transduction. Transduction efficiency was measured after 72 h puromycin treatment using CellTiter-Glo (CTG, Promega) (50 µl of cell suspension + 50 µl CTG), measured with a Fluostar plate reader (BMG Labtech). Luminescence values (after subtracting background signal obtained from the average of wells containing only R10) in puromycin-treated wells at each virus concentration were divided by values of non-puromycin-treated wells. A concentration resulting in 10-20% transduction efficiency was selected to ensure that the majority of the cells receive only one sgRNA.

For the genome-scale screen, > 400 million cells were transduced in 12-well plates. In each well, 3 million cells were suspended in the titrated virus volume achieving 10-20% transduction in a total volume of 1 ml/well topped up with R10 in the presence of 8 µg/ml Polybrene. The plates were centrifuged at room temperature at 800 g for 2 h, after which the virus was washed away. Transduced cells were selected with 0.5 µg/ml puromycin (0.9 µg/ml for K562) for 6 days starting 24 h post-transduction (48 h for NALM6). On day 7 post-transduction (day 8 for NALM6), cells were divided into NK-treated and untreated conditions in T-225 flasks with 120 ml R10 and 60 million target cells, with effector-to-target ratios as listed in Tables S3A-S3B. In some screens, several different effector-to-target ratios were used. The cells were passaged every 2-3 days and cultured for a duration of 4-17 days as listed in Tables S3A-S3B. To maintain sufficient selection pressure, NK cells were added to the cultures 1-2 times during the screens. Approximately 60 million cells were pelleted at the end and at earlier timepoints, frozen in −70 °C, and later thawed for genomic DNA extraction using Blood Maxi Kit (Qiagen).

#### Next generation sequencing

Brunello and Calabrese screens: Preparation of DNA for next generation sequencing was undertaken using a two-step PCR protocol as previously described (Shalem et al., 2014). Briefly, DNA was extracted from frozen cell pellets (3×10^7^ cells; Blood & Cell Culture DNA Maxi Kit, Qiagen) per manufacturer’s instructions. DNA concentration was quantified by UV-spectroscopy (NanoDrop 8000; ThermoFisher Scientific). In the first PCR, sgRNA loci were selectively amplified from a total of 160 µg of genomic DNA (10 µg DNA per sample x 16 reactions, 100 µl volume) using primers described in Table S3C and Phusion^®^ High-Fidelity DNA Polymerase (New England Biolabs, Beverly, MA). This provides approximately 300X coverage for sequencing. A second PCR was performed using 5 µl of the pooled Step 1 PCR product per reaction (1 reaction per 10,000 sgRNAs; 100 µl reaction volume) to attach Illumina adaptors and to barcode samples (Table S3C). Primers for the second PCR included a staggered forward primer (to increase sequencing complexity) and an 8bp barcode on the reverse primer for multiplexing of disparate biological samples (Table S3C). PCR replicates were combined, gel normalized (2% w/v) and pooled, then the entire sample run on a gel for size extraction. The bands containing the amplified and barcoded sgRNA sequences (approximately 350-370 bp) were excised and DNA extracted (QIAquick Gel Extraction Kit, Qiagen). Multiplexed samples were then sequenced at the Molecular Biology Core Facility (Dana-Farber Cancer Institute) and/or The Genomics Platform (Broad Institute) using an Illumina NextSeq 500 (Illumina, San Diego, CA), allowing 4×10^8^ individual reads per multiplexed sample.

GeCKOv2 screens: Amplicons containing sgRNA sequences were amplified with a 2-step PCR protocol using primers flanking the sgRNA cassette (Table S3C) as previously described (Dufva et al., 2020a). Briefly, the following overhangs were added to the locus-specific primers to make them compatible with the index primers: Adapter1 (before locus specific forward primer 5’-3’), Adapter2 (before locus specific reverse primer 5’-3’). The first PCR was performed using 1200 ng of sample DNA and the locus-specific primers, with 96 separate amplifications for each sample. After amplification, all reactions were pooled for the second PCR, in which index primers 1 and 2 and seven identical reactions for each sample pool were used, with a unique combination of dual indexes for each of the sample pools. The seven amplified and indexed reactions were pooled together and purified with Agencourt AMPure XP beads twice. Sample pools were sequenced with Illumina HiSeq 2000 System (Illumina) using read length PE100 or NovaSeq 6000 System (Illumina) using read length PE100.

#### Data analysis

Screen data were analyzed using MAGeCK v0.5.2 and v0.5.7 (Li et al., 2014). Forward direction reads were aligned to the GeCKO v2 library sgRNA sequences using the mageck count function with default parameters. Comparisons across conditions were performed on the resulting sgRNA read count matrix using the mageck test function with default parameters. For the GeCKO v2 screens with the K562, MOLM14, SUDHL4, and NALM6 cells, all NK-treated and untreated samples from different replicates and E:T ratios were respectively pooled together for the MAGeCK test analysis of each cell line. GSEA was run using fgsea (Korotkevich et al., 2021) on gene lists ranked based on signed MAGeCK p values. For comparison of the NALM6 NK cell screen with CAR T cell screen in the same cells, CAR T cell screen data were downloaded from Supplemental Table 5 (Supplemental File 6) in (Dufva et al., 2020a).

### CRISPR screen with focused sgRNA library

Sub-genome scale CRISPR gene editing screens to validate determinants of tumor cell response versus resistance to NK cells in a pooled manner were performed using the same reagents and protocols described in the genome-scale *Brunello* section above. Six hundred thirty-five target genes were selected by pooling top hits and biologically relevant hits from our MM cell screens and our solid tumor screens (Sheffer et al., 2021). Olfactory receptor (OR) genes, which are generally not expressed nor considered to influence tumor cell survival and immune responses, were used to establish a control distribution of sgRNAs. A total of 4,000 sgRNAs targeting screen hits and OR gene control sgRNAs were cloned into lentiCRISPRv2 (a gift from Feng Zhang, Addgene plasmid # 52961), with an additional G added in the beginning of the sgRNA sequence when indicated (Table S3O).

Target cell lines MM.1S, LP1 and KMS11 were co-cultured with donor-derived, IL-2 expanded NK cells (same donor as genome-scale screens) or left untreated, in three biological replicates at the following E:T ratios: LP1 and KMS11 1:2, while MM.1S were treated at 2:1 in one experiment and 1:1 in a subsequent experiment.

After each screen, DNA extraction, PCR amplification, next generation sequencing, and processing of sequencing data were performed as described for genome-scale screens above.

One-sided test for enrichment and depletion of the sgRNAs and sgRNA rank aggregation was performed for each gene using MAGeCK, with default parameter settings. OR genes were used to establish a control distribution of sgRNAs for the rank aggregation procedure. For validation purposes, only those genes included among the top 200 in each genome-scale screen were included in the analysis per each cell line.

### Individual gene CRISPR validations

Single-guide RNAs targeting screen hits and non-targeting control sgRNAs were cloned into lentiCRISPRv2 (a gift from Feng Zhang, Addgene plasmid # 52961), with an additional G added in the beginning of the sgRNA sequence (Table S3P). Lentivirus was produced and luciferase-expressing cells were transduced as described above for the GeCKO library virus. Cells were selected using 0.5 μg/ml puromycin (0.9 μg/ml for K562) prior to experiments.

Cytotoxicity assays using a luciferase readout were performed by plating 10,000 luciferase-expressing target cells harboring each sgRNA were on a 384-well plate alone or with expanded NK cells at 1:2 effector-to-target ratio in a total volume of 25 μl with six replicate wells. Plates were incubated at 37 °C and 5% CO_2_ for 48 h, after which 25 μl ONE-Glo reagent was added to each well luminescence measured with a Pherastar FS plate reader. Raw luminescence values were normalized to the average of technical replicates of target cells carrying each sgRNA cultured without NK cells and average log2 fold changes were calculated between NK-treated and untreated wells for each sgRNA.

### Analysis of CRISPR screen hit mutations and gene expression

Mutations in CRISPR screen hit genes (p < 0.00005 and FDR < 0.2 in any of the screens) were queried from cBioPortal using the following datasets: Chronic Lymphocytic Leukemia (Broad, Nature 2015), Diffuse Large B-Cell Lymphoma (Duke, Cell 2017), Multiple Myeloma (Broad, Cancer Cell 2014), Acute Myeloid Leukemia (TCGA, PanCancer Atlas), Acute Lymphoblastic Leukemia (St Jude, Nat Genet 2016).

Processed gene expression data from normal cell types from BLUEPRINT and ENCODE were downloaded from https://github.com/dviraran/SingleR/blob/master/data/blue_print_encode.rda.

### Multi-omics correlations with PRISM-based NK cell sensitivity

Genetic subtypes of the cell lines were annotated based on previous studies as listed in Table S2A. A data matrix containing genomic and other multi-omic features was generated for systematic pairwise correlation analyses between PRISM AUC and genomic features in CCLE data. CCLE 2021 quartile 4 data was downloaded from https://depmap.org/portal/download/all/. A feature matrix comprising all available data levels was built, harmonizing sample names by DepMap-ID (columns) and categorizing by features (rows) as numeric or binary. Each feature was annotated as NUMERIC|BINARY:DATATYPE:FEATURE using the following abbreviations: GEXP, gene expression; RPPA, protein expression; METH, methylation; CNVR, copy number variation, GNAB, mutation; MIRN, miRNA; LCMS, metabolomics) were distinguished from each other. In all instances, missing data was reported as NA.

Feature pairs were compared using Spearman’s rank correlation followed by p value adjustment using the Benjamini-Hochberg method. In the case of discrete features, only features with at least 5 observations (such as mutations) were used to limit the number of comparisons. Statistical tests were performed to assess whether PRISM-based NK cell sensitivity AUC was correlated with other features, such as gene expression, protein expression, clinical, CNA, mutations, miRNAs, and metabolomics. For genes whose expression correlated with PRISM AUC, the correlation between expression and methylation of the same gene was analyzed. The analyses were performed both across all cell lines and within each cancer type (AML, BCL, B-ALL, T-ALL, MM).

To assess which gene sets were enriched in samples sensitive or resistant to NK cells, GSEA was run using fgsea (Korotkevich et al., 2021) on gene lists ranked based on signed p values of the correlation with PRISM AUC.

Features identified using the pairwise correlation analyses were visualized at the sample level with heatmaps generated using ComplexHeatmap (Gu et al., 2016). The enriched gene sets identified by GSEA were visualized at the sample level using GSVA (Hänzelmann et al., 2013).

### Patient genomic data analysis

#### Data collection and preprocessing

Feature matrices containing clinical data, processed gene expression values, mutations, CNAs, and subtypes of DLBCL patients from Reddy et al. (Reddy et al., 2017), Chapuy et al. (Chapuy et al., 2018) and the TCGA dataset; MM patients from the CoMMpass dataset (Manojlovic et al., 2017); and AML patients from the TCGA dataset (2013) preprocessed as previously described (Dufva et al., 2020b) were downloaded from Synapse (Reddy et al. DLBCL: https://www.synapse.org/#!Synapse:syn21995529, Chapuy et al. DLBCL: https://www.synapse.org/#!Synapse:syn21991358, TCGA DLBCL: https://www.synapse.org/#!Synapse:syn21995730, CoMMpass MM: https://www.synapse.org/#!Synapse:syn21995455, TCGA AML: https://www.synapse.org/#!Synapse:syn21995719. Clinical data, processed gene expression values, mutations, and subtypes of 262 T-ALL patients were downloaded from supplementary tables 1, 5, 8, and 15, respectively (Liu et al., 2017).

Processed methylation beta values of 109 T-ALL patients and 20 samples of normal thymocytes were downloaded from GEO (GSE155333). Genetic subtypes were obtained from supplementary table 6 (Roels et al., 2020).

#### Pairwise correlation analysis and visualization

Data matrices containing genomic and other multi-omic features as well as clinical annotations were generated as described above for systematic pairwise correlation analyses, including correlations with the NK cell sensitivity signatures (Figure 5) and with expression of CRISPR screen hits in patient data (Figures 4 and S4). To find patient samples with similar molecular phenotypes as the NK-sensitive cell lines, NK cell sensitivity signatures were obtained by taking 50 genes most significantly correlating with sensitivity to NK cells based on PRISM AUC. The 50 genes were used to calculate an enrichment score of the NK sensitivity signature for each patient sample using GSVA. The NK cell sensitivity signatures were derived separately from MM, T-ALL, and BCL cell lines for use in the corresponding patient datasets. Spearman’s rank correlation followed by p value adjustment using the Benjamini-Hochberg method was used to assess whether the NK cell sensitivity signatures were correlated with other features, such as gene expression, clinical, CNAs, or mutations. A similar approach was used to test if the expression of CRISPR screen hits correlated with methylation or copy number of the same gene.

To visualize identified associations of CRISPR/PRISM features with patient genomic and clinical data as dot plots, differential expression between a sample group and all other samples was calculated using Wilcoxon rank sum test. For UMAP visualizations, expression values of 15% of the most variable genes were used for dimensionality reduction using the umap R package (McInnes et al., 2020).

### Single-cell transcriptomics CRISPR screens

#### Experiments and preparation of scRNA-seq libraries

To generate lentiviral sgRNA libraries for single-cell CRISPR screens, guides targeting screen hits (three sgRNAs for each gene) and non-targeting control sgRNAs (four for K562, six for other screens; Table S6A) were cloned into CROPseq-Guide-Puro (a gift from Christoph Bock, Addgene # 86708) (Datlinger et al., 2017) or into CROP-sgRNA-MS2 (a gift from Wolf Reik, Addgene # 153457) (Alda-Catalinas et al., 2020). Lentivirus was produced and cells were transduced as described above using different concentrations of virus. Cells were selected with puromycin (0.5 µg/ml for K562, SUDHL4, and NALM6; 1 µg/ml for MM1.S, LP1, and KMS11) and cells transduced with a concentration resulting in a 10-20% transduction efficiency based on a viability assay described above were selected for screening. Target cells were co-cultured with day 17 feeder cell-expanded NK cells for 24 h at 1:16 and 1:4 effector-to-target ratios (only 1:16 for K562) or left untreated, and both conditions were subjected to scRNA-seq after washing twice with 10 ml PBS + 0.04% BSA.

The Chromium Single Cell 3’ RNAseq run and library preparation were done using the 10x Genomics Chromium Next GEM Single Cell 3’ Gene Expression v3.0 chemistry (K562), v3.1 chemistry (SUDHL4, NALM6, MM1S), or v3.1 Dual Index chemistry (LP1, MM1S CRISPRa). CROP-seq guide sequencing libraries were prepared using nested PCRs described in Hill et al. 2018 and https://github.com/shendurelab/single-cell-ko-screens#enrichment-pcr. Briefly, 13 ng of full length 10x cDNA was used as template for the first round of amplification. The subsequent 2nd and 3rd PCR reactions were done using SPRIselect Reagent (1.0X) purified and 1:25 diluted PCR product as template. Optimal amplification cycles were selected based on quantitative PCR analysis. The guide sequencing libraries were sequenced alongside the 3’ GEX libraries with approximately 10% read depth when compared to 3’ GEX libraries. The K562, SUDHL4, NALM6, and MM1S sample libraries were sequenced on Illumina NovaSeq 6000 system using the following read lengths: 28bp (Read 1), 8bp (i7 Index), 0 bp (i5 Index) and 89bp (Read 2). The LP1 and MM1S CRISPRa sample libraries were sequenced on Illumina NovaSeq 6000 system using the following read lengths: 28bp (Read 1), 10bp (i7 Index), 10bp (i5 Index) and 90bp (Read 2).

#### Data analysis

Data preprocessing was performed using 10x Genomics Cell Ranger v3.1 (K562, SUDHL4, NALM6, MM1S) or v6.0.2 (LP1, MM1S CRISPRa) pipelines. The ‘cellranger mkfastq’ function was used to produce FASTQ files and ‘cellranger count’ to perform alignment, filtering, and UMI counting. The Illumina bcl2fastq v2.2.0 was used to run mkfastq function and alignment was done against the human genome GRCh38.

FASTQ files of the targeted sgRNA amplification libraries were run through Cell Ranger count v3.1.0 pipeline. UMI counts of guides associated with each cell were extracted using the get_barcodes.py script downloaded from https://github.com/shendurelab/single-cell-ko-screens (Hill et al., 2018). To assign guides to cells, cells harboring sequences with > 10 UMI counts and accounting for > 50% of the UMI counts in the cell were included in the analysis. Out of these, cells in which the second most frequent guide accounted for > 20% of the UMI counts were considered to express two guides and were removed from the analysis.

The R package Seurat (v4.0.4) (Stuart et al., 2019) was used for further scRNA-seq data processing. Cells with > 10-15% mitochondrial gene counts, > 50% or < 5% ribosomal gene transcripts, < 3,000 UMI counts, or < 300 or > 10,000 detected genes were filtered out. After log-normalization, the highly variable genes were calculated with the FindVariableFeatures function in Seurat using the ‘vst’ selection method. Data were scaled, clusters were defined based on PCs with a standard deviation > 2 using the FindNeighbors and FindClusters functions, and cell types were annotated using SingleR. Clusters comprising NK cells, doublets, or low-quality cells were removed. The sgRNA–cell assignments were merged with the expression object, which was subsetted to cells assigned a single sgRNA. Differential expression between cells expressing guides targeting a gene and non-targeting controls or between untreated or NK cell-treated control sgRNA-carrying cells was performed with a Student’s *t*-test using the FindMarkers function in Seurat with logfc.threshold = 0.1. Multiple testing correction using the Bonferroni method was performed separately for each perturbation. Similarity of the differential expression gene lists across perturbations was assessed using the CompareLists function in the OrderedList package (v1.64.0).

The mixscape tool in Seurat was used to detect perturbations with a transcriptomic phenotype and visualize their relative differences as previously described (Papalexi et al., 2021). CalcPerturbSig was used to calculate perturbation signatures reflecting the perturbation-specific differences between cells expressing gene-targeting guides and cells expressing control guides, and cells were classified as perturbed or non-perturbed using RunMixscape with logfc.threshold = 0.025 and gene.count = 5. Cells classified as non-perturbed were removed from the analysis and the similarity of the perturbations was visualized as a UMAP based on linear discriminant analysis computed using the MixscapeLDA function.

The core NK cell response score was calculated as described above and differential enrichment of the score in various perturbations was calculated with the Student’s *t*-test using the FindMarkers function in Seurat similarly as for genes.

#### Comparison with patient data

For the genes perturbed in CROP-seq experiments mutated in at least 5 patients with either MM (CoMMpass), DLBCL (Reddy et al.), or AML (TCGA), differentially expressed genes between patients with and without mutations were determined using limma (Ritchie et al., 2015). Genes significantly differentially expressed in the same direction at a nominal p value threshold of 0.05 in both CROP-seq and in patients when the same gene was knocked out or mutated, respectively, were identified.

## DATA AVAILABILITY

The results of the study can be **interactively explored** at: https://immunogenomics.shinyapps.io/nkheme/. All processed and raw sequencing data generated in this study will be made publicly available upon final publication of this work.

## REAGENTS AND RESOURCES

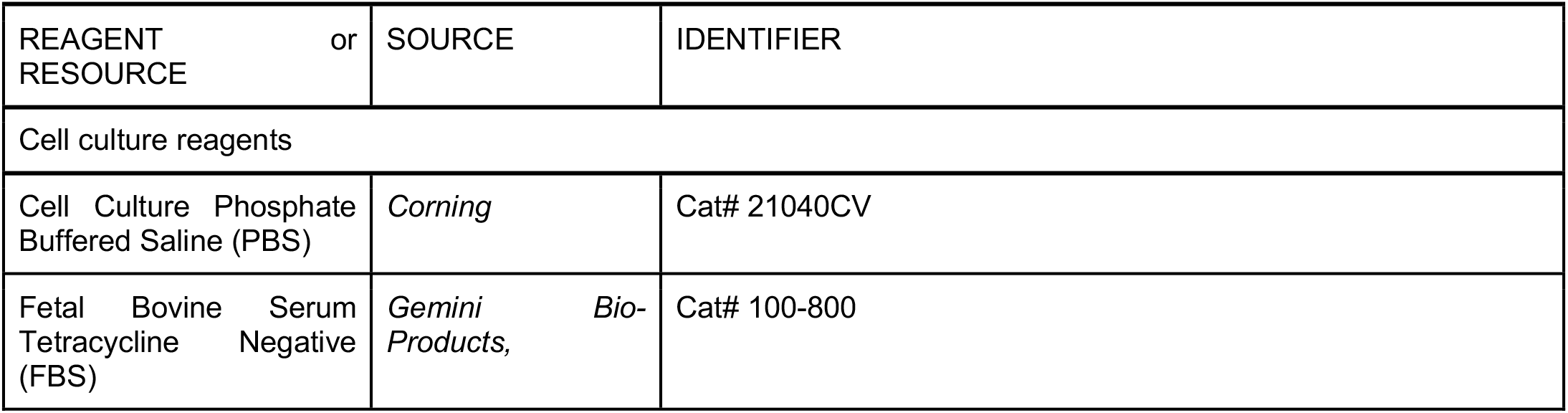
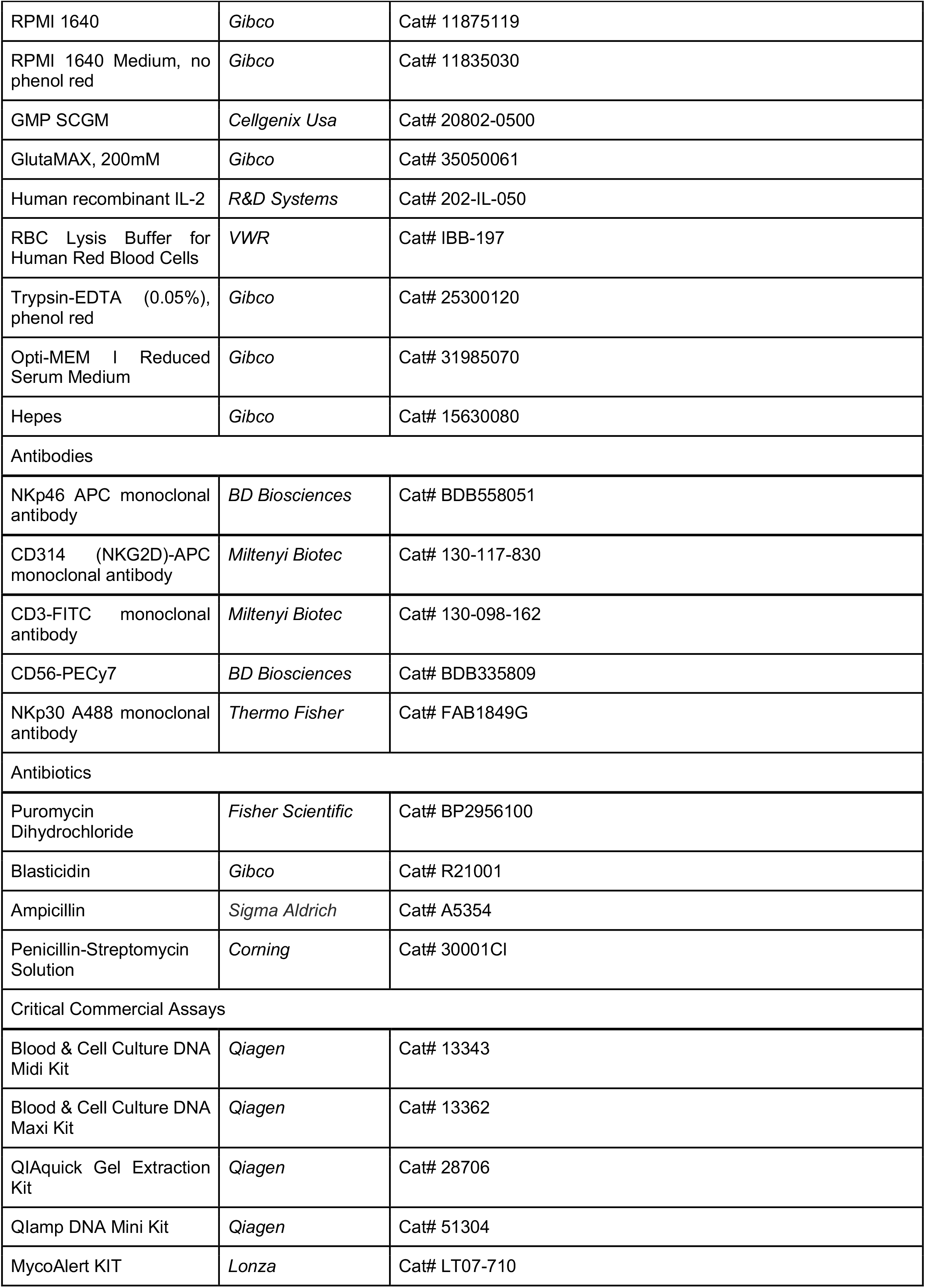
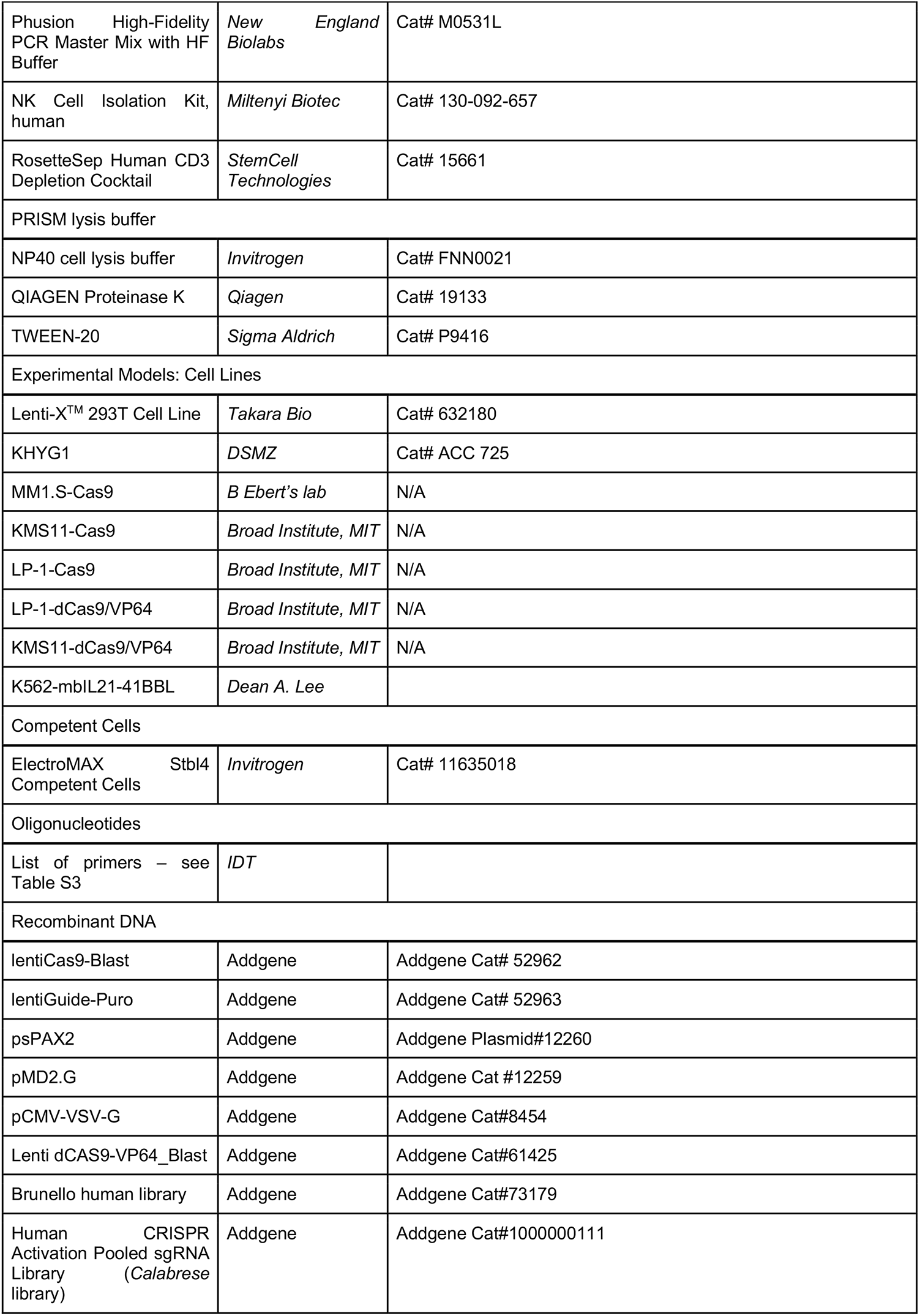
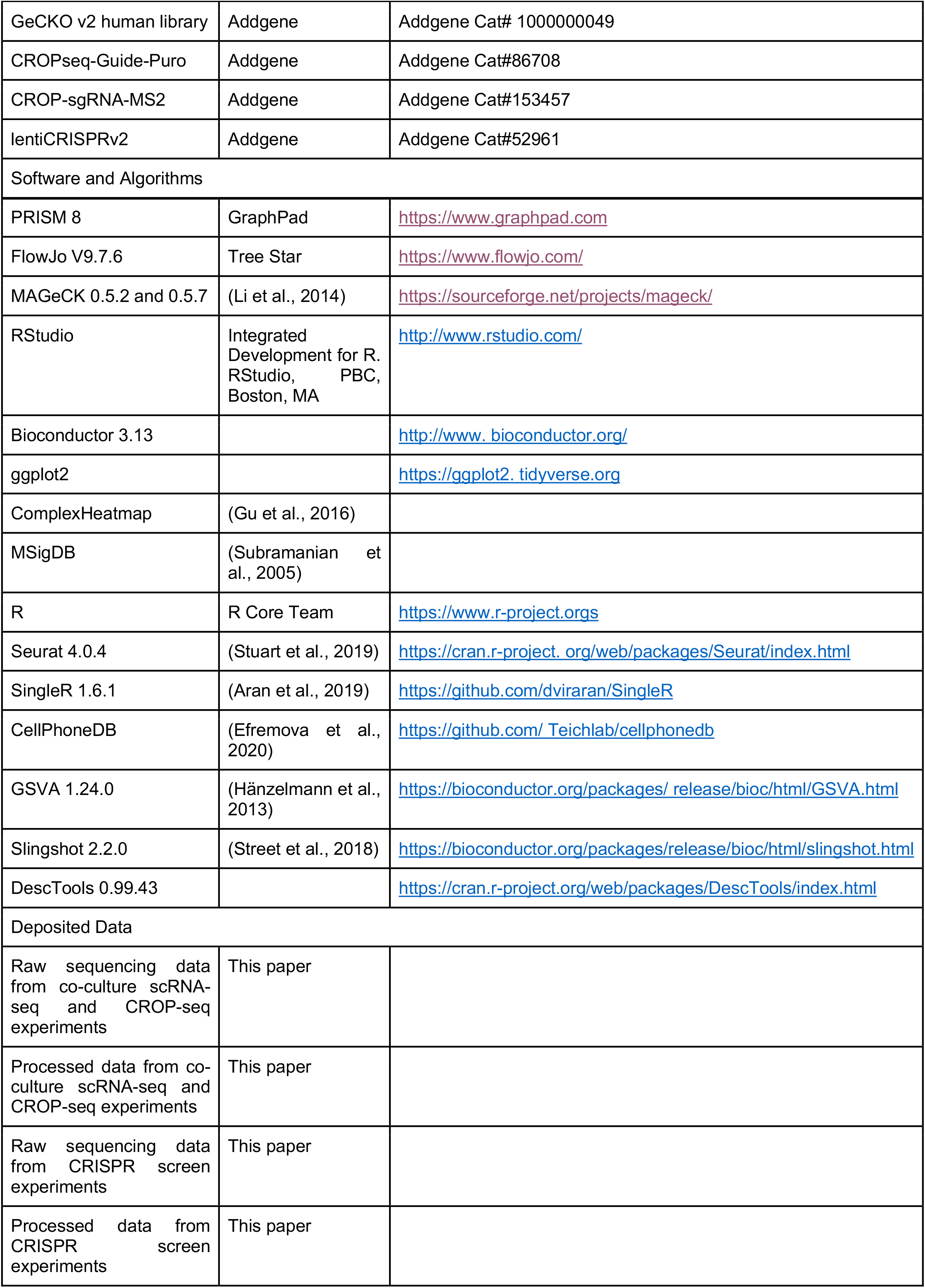
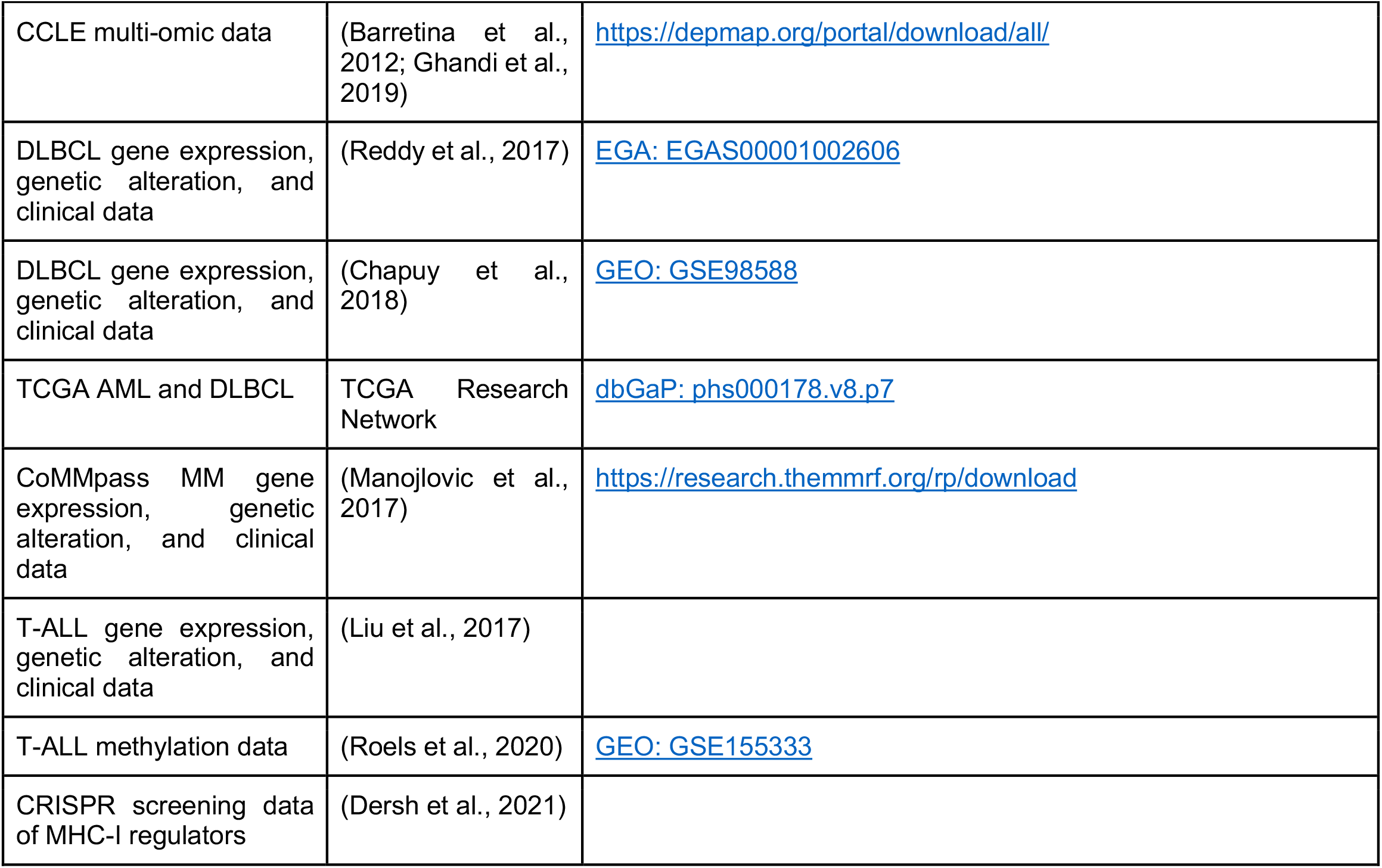

## ACKNOWLEDGEMENTS

The authors would like to thank Prof. Yenan Bryceson (Karolinska Institute, Sweden) and Drs. Mikko Keränen, Oscar Brück, Mikko Myllymäki, Sofie Lundgren, and Leo Meriranta for insightful comments on this manuscript. The Biomedicum Sequencing laboratory and Virus core supported by the Biocentrum Helsinki and Helsinki Institute of Life Science (HiLIFE) as well as the The Biomedicum Helsinki Flow Cytometry Core Unit are acknowledged for their services. Single cell sequencing was performed at FIMM Single-Cell Analytics and Sequencing units supported by HiLIFE and Biocenter Finland. The primary NK cells were extracted from buffy coats provided by the Finnish Red Cross Blood Service biobank.

The study was supported by grants from the Cancer Foundation Finland, Academy of Finland, Sigrid Juselius Foundation, Gyllenberg Foundation, State funding for the University-level Health Research in Finland, and HiLIFE fellow funds (to S.M.).This work was also supported by the Stand Up To Cancer (SU2C) Convergence 2.0 Grant (S.G., M.S., C.S.M.), a collaboration by SU2C and the Society for Immunotherapy of Cancer; the SU2C Phillip A. Sharp Award for Innovation in Collaboration (M.S., C.S.M.); the Leukemia and Lymphoma Society (LLS) Scholar Award (C.S.M.); de Gunzburg Myeloma Research Fund (C.S.M.); International Myeloma Society (C.S.M.), the Shawna Ashlee Corman Investigatorship in Multiple Myeloma Research (C.S.M.); Cobb Family Myeloma Research Fund (S.G., C.S.M.); and the Ludwig Center at Harvard (C.S.M.). The collaborating laboratories have also been partially supported by grants NIH R01 CA050947 (C.S.M.), CA196664 (C.S.M.), U01 CA225730 (C.S.M.), the Lauri Strauss Leukemia Foundation (to R.S.), the LLS Translational Research Program (C.S.M.), the Multiple Myeloma Research Foundation (MMRF) Answer Fund (C.S.M.). O.D. was supported by grants from the Biomedicum Helsinki Foundation, Finnish Medical Society, Cancer Foundation Finland, K. Albin Johansson Foundation, Emil Aaltonen Foundation, Ida Montin Foundation, Finnish Blood Disease Research Foundation, and Finnish Hematology Association. S.G. was also supported by the ASCO Conquer Cancer Young Investigator Award, the International Myeloma Society Career Development Award, the Associazione Italiana per la Ricerca sul Cancro (AIRC, Italian Association for Cancer Research).

