## Supplemental figures for "Single-cell functional genomics of natural killer cell evasion in blood cancers"

[1]Hematology Research Unit Helsinki, Helsinki University Hospital Comprehensive Cancer Center, Helsinki, Finland
[2]Translational Immunology Research Program and Department of Clinical Chemistry and Hematology, University of Helsinki, Helsinki, Finland
[3]iCAN Digital Precision Cancer Medicine Flagship, Helsinki, Finland
[4]Department of Medical Oncology, Dana-Farber Cancer Institute, Harvard Medical School, Boston, MA, USA
[5]Broad Institute of MIT and Harvard, Cambridge, MA, USA
[6]Department of Medicine, Harvard Medical School, Boston, MA, USA
[7]Ludwig Center, Harvard Medical School, Boston,MA, USA
[8]Institute for Molecular Medicine Finland (FIMM), HiLIFE, University of Helsinki, Helsinki, Finland
[9]Faculty of Health Sciences, A.I. Virtanen Institute for Molecular Sciences, University of Eastern Finland, Kuopio, Finland
[10]Department of Pathology, St. Jude Children's Research Hospital, Memphis, TN, USA
[11]Department of Computer Science, Aalto University, Espoo, Finland
[12]Hematology/Oncology/BMT, Center for Childhood Cancer and Blood Diseases, Nationwide Children's Hospital, Columbus, OH

#### SUPPLEMENTAL FIGURES

Supplemental Figure 1

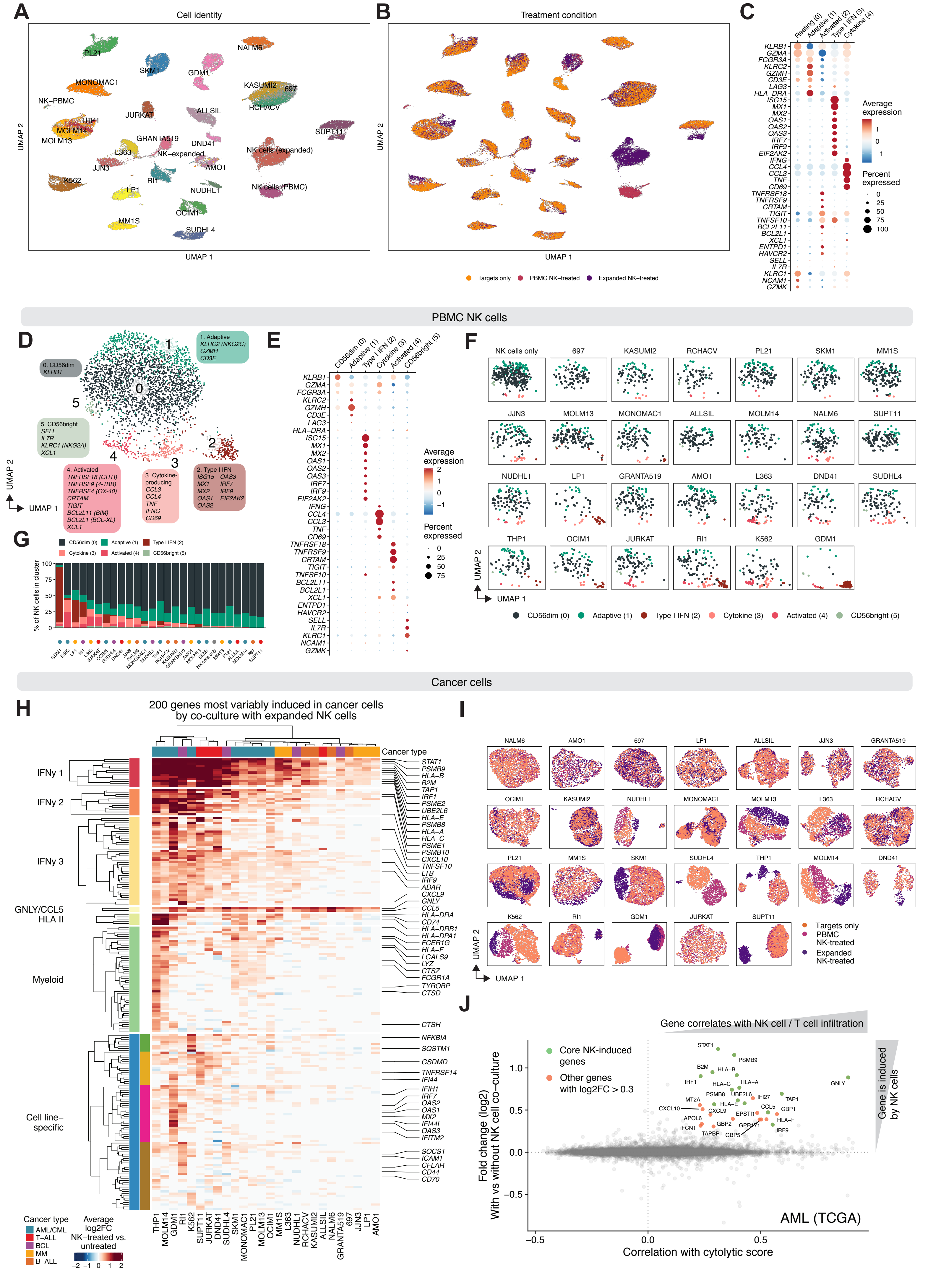

**Figure S1. Multiplexed scRNA-seq of interacting NK cells and blood cancer cells. Related to Figures 1 and 2.**

(A) UMAP visualization of all cells in the co-culture experiments after quality control (n = 61,715). Cells are colored based on cell line/NK cell identity.
(B) As in A, but cells are colored based on culture condition.
(C) Dot plot of expanded NK cell cluster marker genes selected from the genes significantly overexpressed in each cluster compared to other clusters. Color indicates average expression of the gene and dot size indicates percent of cells in which expression of the gene is detected.
(D) UMAP visualization of PBMC NK cells (non-expanded) from all conditions, including co-culture with 26 cell lines and NK cell cultured alone. Cells are colored based on the clusters and marker genes are shown for each cluster selected from the genes significantly overexpressed in each cluster compared to other clusters.
(E) Dot plot of PBMC NK cell cluster marker genes selected from the genes significantly overexpressed in each cluster compared to other clusters. Color indicates average expression of the gene and dot size indicates percent of cells in which expression of the gene is detected.
(F) UMAP visualizations of PBMC NK cells co-cultured with each of the 26 cell lines and NK cells cultured alone. Cells are colored according to clusters.
(G) Bar plot of percentages of PBMC NK cells belonging to different clusters in each condition, including co-culture with 26 cell lines and NK cells cultured alone. Conditions are ordered by the combined percentage of cells in CD56dim and adaptive-like clusters (clusters 0 and 1) in increasing order.
(H) Heatmap of top 200 genes most variably induced across the blood cancer cell lines upon co-culture with expanded NK cells. Cell lines and genes are hierarchically clustered using Euclidean distance and ward.D2 linkage. Gene clusters are annotated based on genes included in the clusters. The ‘Cell-line specific’ cluster is further clustered into four subclusters. Selected genes are labeled on the right. The cancer type of the cell lines is indicated above.
(I) UMAP visualizations of cancer cells cultured alone or with expanded or PBMC-derived NK cells. Cells are colored according to the co-culture condition.
(J) Scatter plot comparing genes induced by NK cell co-culture in target cells and genes correlating with NK and T cell infiltration (cytolytic score) in AML patient samples from TCGA. Genes with significant correlation and differential expression (FDR < 0.05) and scRNA-seq log2 fold change > 0.3 are labeled in red. Genes included in the core NK-induced genes are labeled in green.

### Supplemental Figure 2

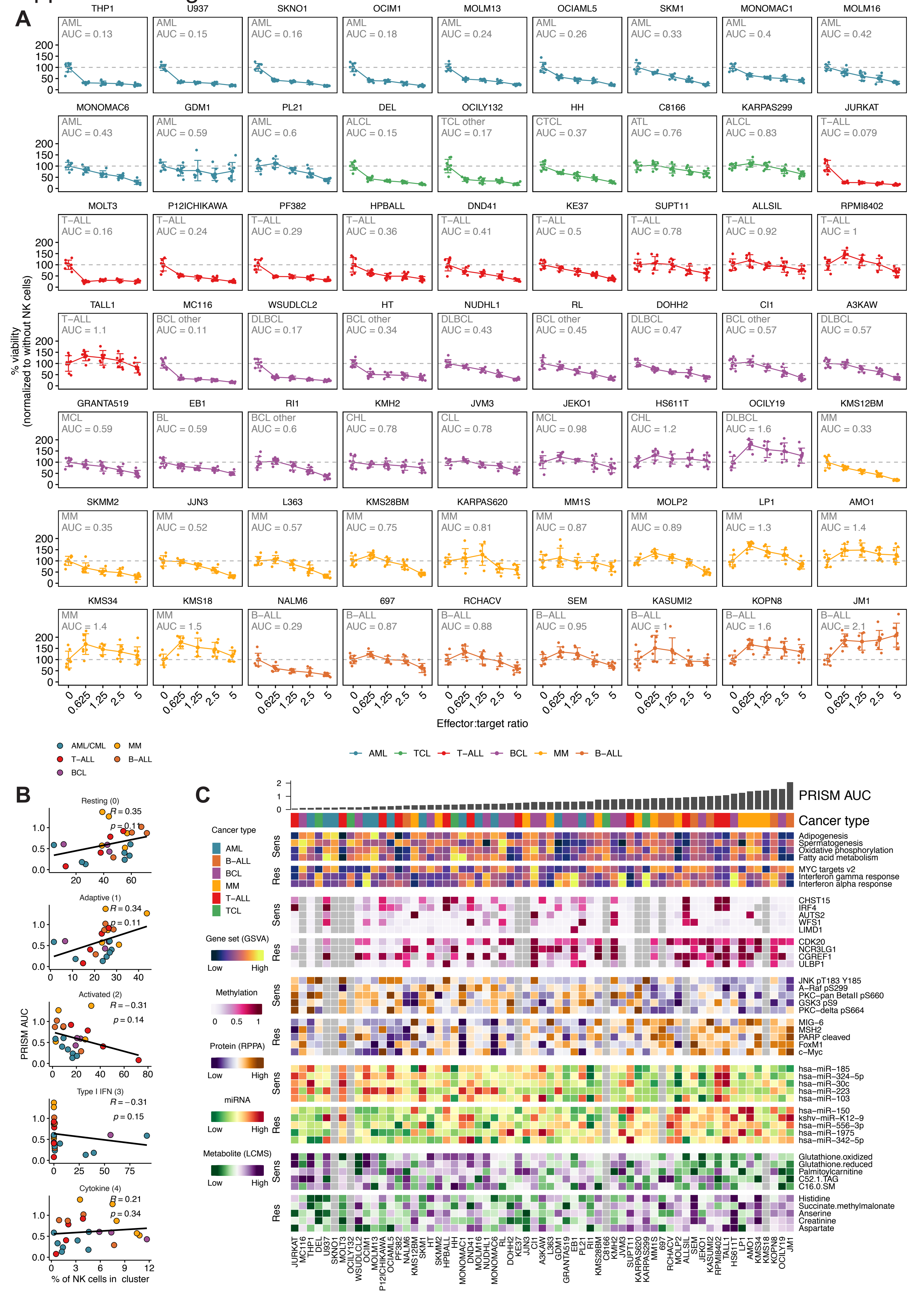

**Figure S2. Sensitivity of individual cell lines to NK cells and multi-omic correlates of NK cell sensitivity. Related to Figure 3.**

(A) Dose-response curves of sensitivity of hematological cell lines to NK cells at different effector-to-target ratios. Cell lines are ordered by cancer type and increasing AUC (decreasing sensitivity) of the dose-response curves. The detailed cancer type and AUC is shown for each cell line. Dots represent technical replicates (n = 8), error bars indicate the standard deviation, and larger dots indicate the mean.

(B) Scatter plots showing the correlation of PRISM AUC (NK cell sensitivity) of cell lines with the fraction of NK cells in different scRNA-seq clusters (Figure 1C) when co-cultured with the corresponding cell lines. Correlation coefficients and p values are obtained using Spearman's correlation.

(C) Heatmap of multi-omic correlates of sensitivity to NK cells across blood cancer cell lines. Cell lines are ordered by sensitivity (PRISM AUC). Hallmark gene sets significantly enriched in resistant or sensitive cell lines, methylation levels of top five genes from figure 5E whose expression most correlates with methylation, and five top protein (RPPA), miRNA, and metabolite features correlating with resistance or sensitivity are shown. All features are scaled except methylation which is shown as beta values.

Supplementary Figure 3

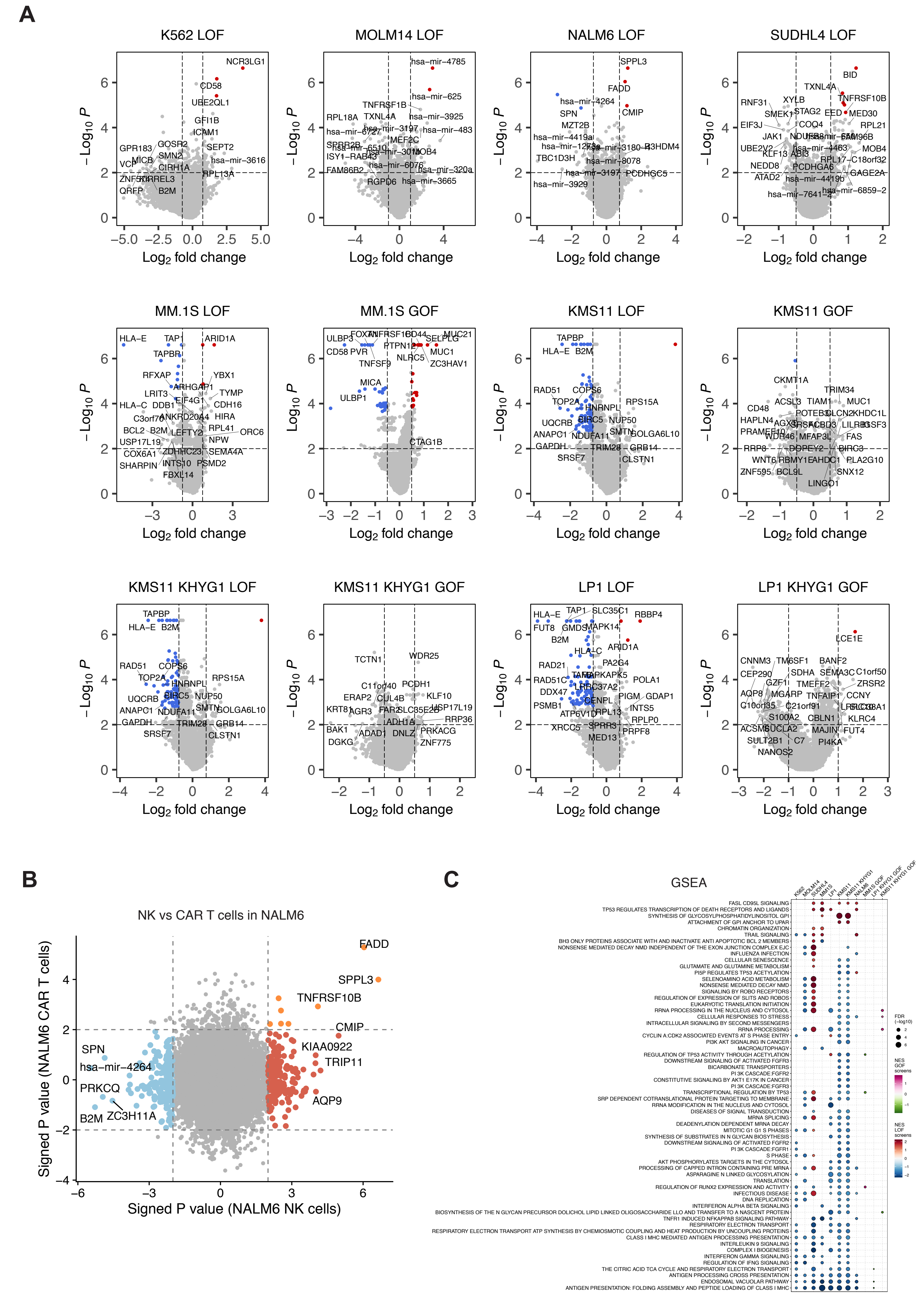

**Figure S3. Hits in individual genome-scale CRISPR screens and gene set enrichment analysis. Related to Figure 4.**

(A) Volcano plots of genome-scale CRISPR screens, showing on the x-axis the average log2 fold change between NK-treated and untreated cells for each gene, while the -log10 p value is shown on the y-axis. Labeled are the top genes that met the significance criteria for log2 fold change > |0.75| and p < 0.01. In red are genes whose sgRNAs were enriched upon exposure to NK cells with FDR < 0.1 and in blue are genes whose sgRNAs were depleted with FDR < 0.1.

(B) Scatter plot comparing NALM6 co-culture screens performed using NK cells or CAR T cells. Genes with negative log2 signed p value in the NK cell screen > 2 or < -2 are labeled red and blue, respectively. Genes with negative log2 signed p value both in the NK cell and the CAR T cell screens > 2 are labeled yellow (genes promoting both NK and CAR T cell cytotoxicity).

(C) Dot plot of gene set enrichment analysis of genes conferring resistance or sensitivity to NK cells in genome-scale CRISPR screens. Shown are Reactome gene sets recurrently enriched in at least two screens. Color indicates normalized enrichment score and dot size indicates the negative log10 p value, with only dots where p < 0.05 shown.

### Supplemental Figure 4

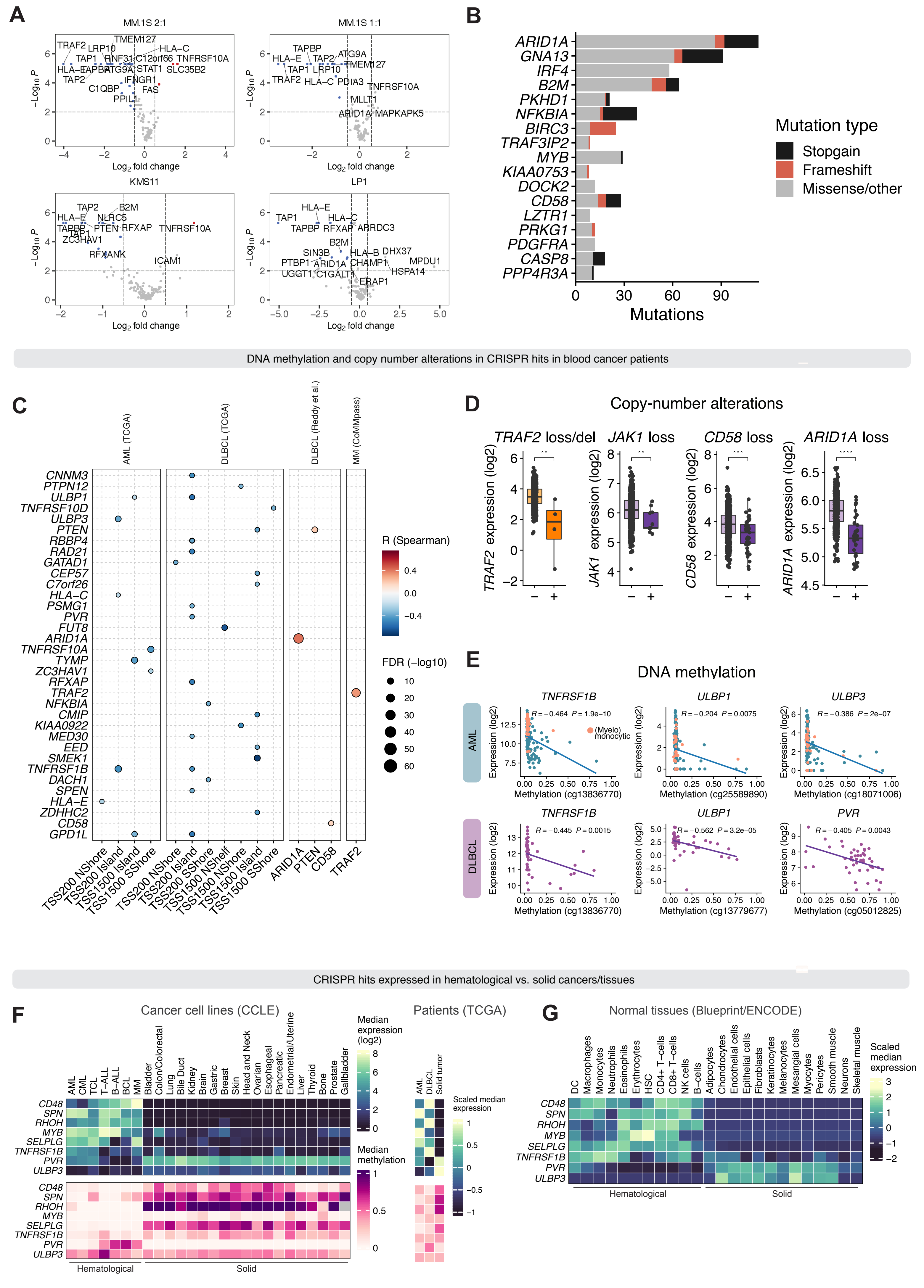

**Figure S4. Focused library validations of CRISPR screens and genetic alterations and transcriptional regulation of screen hits in cancer patients. Related to Figure 4.**

(A) Volcano plots of focused library CRISPR screens, showing on the x axis the average log2 fold change between NK-treated and untreated cells for each gene, while the -log10 p value is shown on the y axis. In red are genes whose LOF decreased response to NK cells and had an FDR < 0.1, in blue are genes whose LOF increased response to NK cell attack and had an FDR < 0.1.

(B) Mutations in CRISPR screen hits in patients with hematological malignancies. Bar plot shows the total number of mutations in each gene across the cohorts shown in Figure 4F, with color indicating the mutation type.

(C) Dot plot of correlation of expression of CRISPR screen hits with DNA methylation and copy number alterations in patient datasets. Shown are only CRISPR screen hit genes whose expression correlates with either methylation of copy number in one of the datasets.

(D) Expression of selected CRISPR screen hit genes whose expression correlates with copy number. '+' indicates altered samples and '-' non-altered samples. Color indicates cancer type. P values are obtained using a Wilcoxon rank sum test.

(E) Scatter plots comparing expression of selected CRISPR screen hit genes with DNA methylation. The methylation array probe is indicated in parentheses. P values and correlation coefficients are obtained using Spearman's correlation. Orange color indicates AML samples with myelomonocytic (FAB M4) or monocytic (FAB M5) differentiation.

(F) Heatmaps showing median expression (top) and methylation (bottom) of selected CRISPR screen hits preferentially expressed in blood cancers or solid tumors in the indicated blood cancer subtypes and solid tumors in CCLE cell lines (left) and TCGA patient samples (right).

(G) Heatmap showing median expression of selected CRISPR screen hits preferentially expressed in blood cancers or solid tumors in normal tissues in Blueprint/ENCODE data.

### Supplemental Figure 5

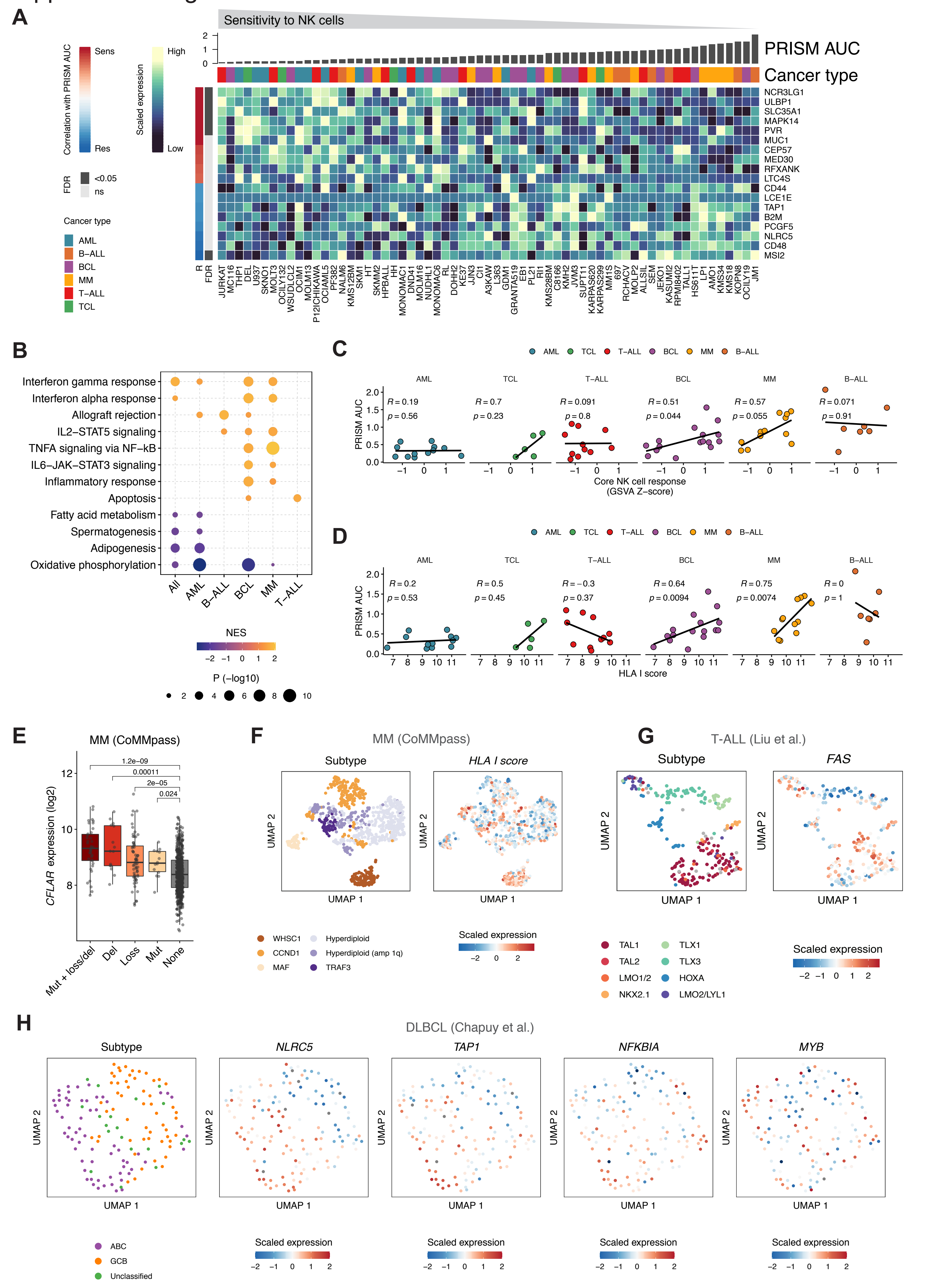

**Figure S5. Molecular correlates of NK cell sensitivity in cell lines and integration with patient genomic data. Related to Figure 5.**

(A) Heatmap of CRISPR screen hits whose expression correlates with sensitivity to NK cells. Cell lines are ordered by sensitivity (PRISM AUC). Only genes with Spearman's correlation $p < 0.05$ are shown. FDR is indicated on the left.

(B) Dot plot of gene sets recurrently correlated with NK cell sensitivity in CCLE data. Gene sets correlated in at least two categories, either two cancer types or one cancer type and across all cancers, are shown. Color indicates GSEA normalized enrichment score (NES). Dot size indicates the negative log10 p value, with only dots where $p < 0.05$ shown.

(C) Scatter plots comparing expression of the core NK cell response gene set as GSVA score with NK cell sensitivity (PRISM AUC) stratified by cancer type. Correlation coefficients and p values are obtained using Spearman's rank correlation.

(D) Scatter plots comparing HLA I score summarizing expression of HLA I complex genes with NK cell sensitivity (PRISM AUC) stratified by cancer type. Correlation coefficient and p value are obtained using Spearman's rank correlation.

(E) Box plot of *CFLAR* expression in MM transcriptomic data from CoMMpass (n = 767) stratified by *TRAF3* alterations. P values are obtained using a Wilcoxon rank sum test. Boxes indicate IQR with a line at the median. Whiskers represent the min and max values at most 1.5 IQR from the quartiles.

(F) UMAP of MM transcriptomic data from CoMMpass (n = 767) as in Figure 5C with molecular subtypes and HLA I score colored on the plot.

(G) UMAP of T-ALL transcriptomic data from Liu et al. (n = 262) as in Figure 5F with molecular subtypes and *FAS* expression colored on the plot.

(H) UMAPs of DLBCL transcriptomic data from Chapuy et al. (n = 137) as in Figure 5I with molecular subtypes and *NLRC5*, *TAP1*, *NFKBIA*, and *MYB* expression colored on the plots.

### Supplementary Figure 6

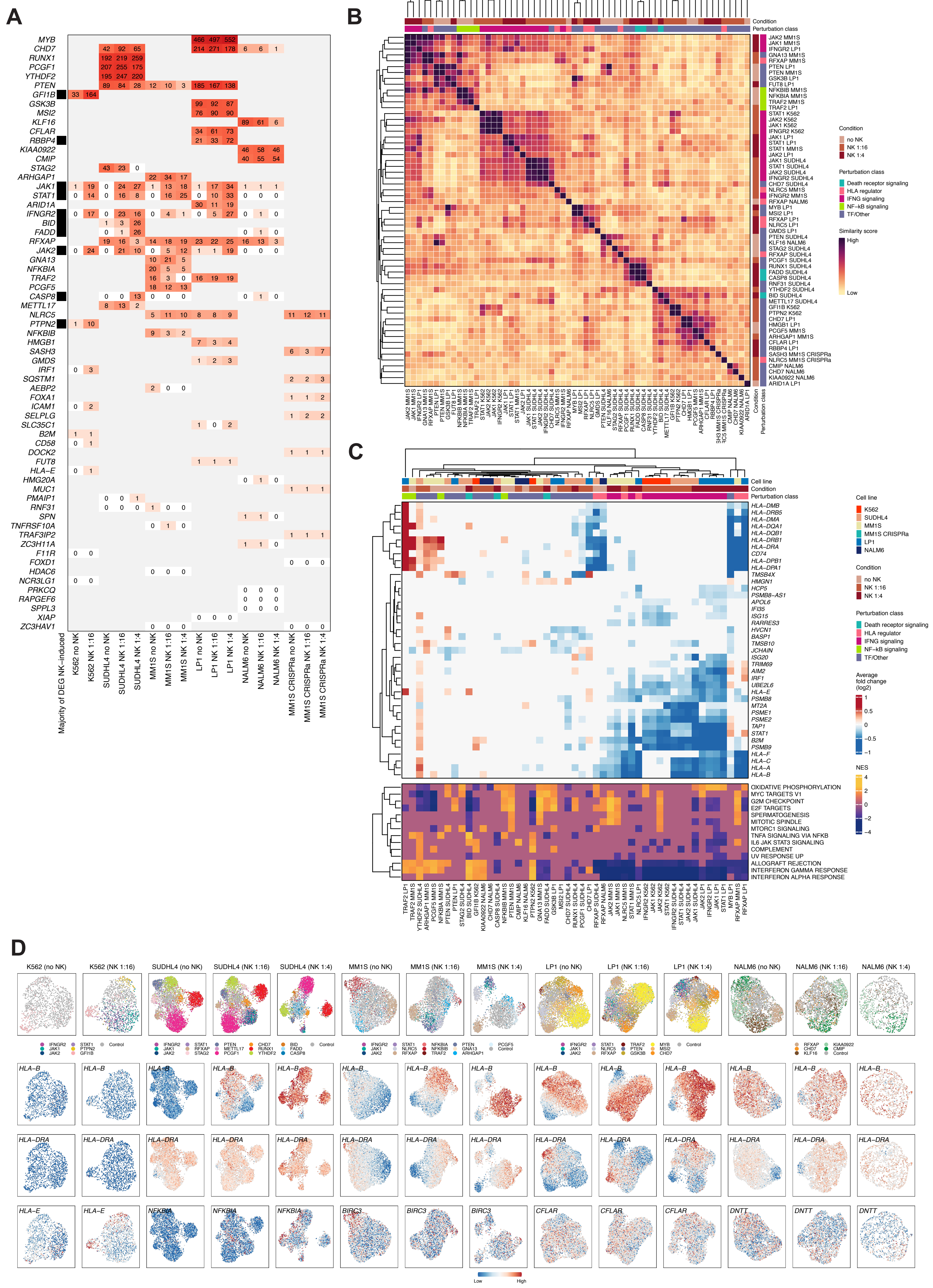

**Figure S6. Comparison of global transcriptomic effects across perturbations in single-cell transcriptomics CRISPR screens. Related to Figure 6.**

(A) Heatmap showing numbers of differentially expressed genes (DEGs) with each perturbation (y axis) in each cell line and condition (x axis). Darker red shades indicate higher number of DEGs and grey rectangles indicate perturbations not included in the cell line. Perturbations in which the majority of DEGs are found in the NK-treated conditions are indicated with a black square on the left.

(B) Heatmap of pairwise similarities of DEGs between different CROP-seq perturbations. Similarity scores were determined by the OrderedList package. Rows and columns are clustered using Euclidean distance and complete linkage. Conditions and functional classes of the perturbations are shown at the top and at the right. For each perturbation, the condition with the highest number of DEGs is shown. Only perturbations with at least five differentially expressed genes are included in the analysis.

(C) Heatmap of genes and pathways recurrently differentially expressed across the different CROP-seq perturbations. Genes upregulated or downregulated and Hallmark pathways enriched or depleted in over five perturbations in the same direction are shown. Rows and columns are clustered using Euclidean distance and complete linkage. Cell lines, conditions, and functional classes of the perturbations are shown at the top. For each perturbation, the condition with the highest number of DEGs is shown. Only perturbations with at least five differentially expressed genes are included in the analysis.

(D) UMAP visualizations of CROP-seq data after running linear discriminant analysis (LDA) using mixscape in the indicated cell lines in different conditions. Perturbed genes are colored on the plots in the top row, and examples of differentially expressed genes are colored in the two bottom rows. Only cells classified as knockout or non-targeting (‘Control cells’ colored gray) by mixscape are shown. For the K562 ‘no NK’ condition, the UMAP is computed using principal components of the highly variable genes instead of LDA due to only one perturbation having a significant phenotype.

### Supplementary Figure 7

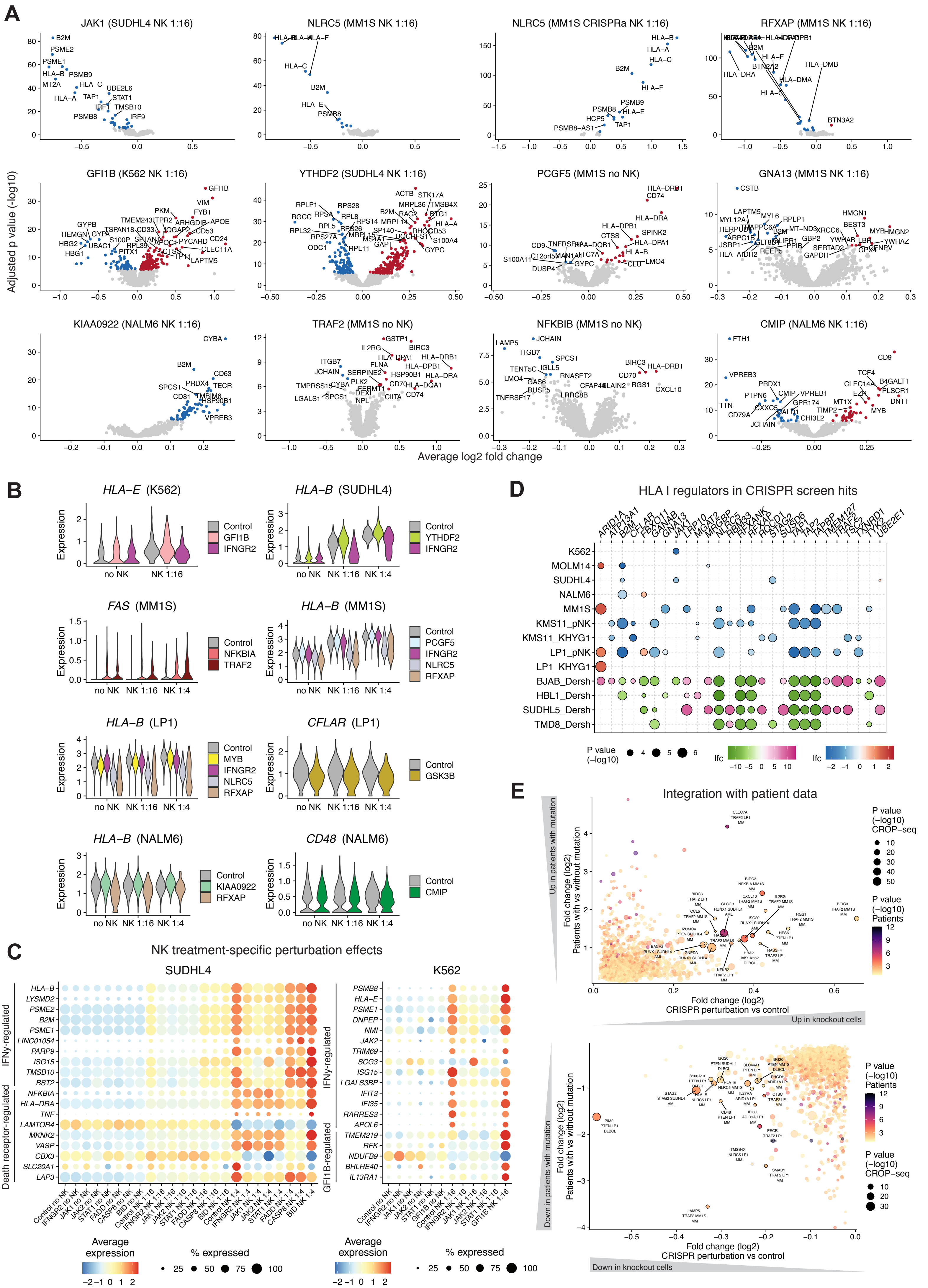

**Figure S7. Transcriptomic effects of individual perturbations in single-cell transcriptomics CRISPR screens. Related to Figure 6.**

(A) Volcano plots showing differentially expressed genes of selected perturbations compared to control (non-targeting sgRNAs). The perturbed gene, cell line, and condition are indicated above the plots. Genes differentially expressed with adjusted p value < 0.05 are labeled red (positive log2 fold change) or blue (negative log2 fold change). Genes with adjusted p value >= 0.05 are labeled gray.

(B) Violin plots of selected genes and perturbations in each treatment condition (no NK, NK 1:16, NK 1:4).

(C) Dot plots showing expression of top DEGs only found under NK cell-treatment for SUDHL4 and K562 cells. Shown are perturbations (x axis) with the highest fraction of DEGs only found under NK cell-treatment and top DEGs (y axis) for each perturbation (top 3 for SUDHL4, top 5 for K562). Groups of DEGs related to IFNγ pathway perturbations (*IFNGR2*, *JAK1*, *JAK2*, *STAT1*), death receptor pathway perturbations (*FADD*, *CASP8*, *BID*), and GFI1B perturbation are indicated on the left side of the plots. Dot color indicates scaled average expression of the gene across cells harboring the perturbation and dot size indicates the percentage of cells in which expression of the gene is detected.

(D) Dot plot showing the integration of our genome-scale CRISPR screen data and data published by Dersh et al., comparing DLBCL cell lines with high levels of expression of HLA-I to HLA-I low. Only genes with p < 0.01 in both sets of screens, in either direction (i.e. enrichment or depletion) are shown. Dot color indicates scaled average log2 fold change of the cells carrying the perturbation compared to control; dot size indicates the negative log10 p value, with only dots where p < 0.01 shown. In the CRISPR dataset, enriched sgRNAs are depicted in red, while depleted sgRNAs are in shades of blue. In the Dersh dataset, negative regulators of class I HLA are depicted in shades of pink, while positive regulators are depicted in shades of green.

(E) Comparison of transcriptomic effects of single-cell CRISPR perturbations and patients with mutations in corresponding genes. Differential expression between patients with or without mutations in CRISPR screen hit genes (y axis) is compared with differential expression between cells where the same gene is perturbed in single-cell CRISPR screens compared to control cells. Only pairs where both differential expressions are significant (p < 0.05) and fold changes are positive (upper plot, upregulated genes) or negative (bottom plot, downregulated genes) are shown. Differentially expressed gene, cell line and perturbation, and cancer type of the patient dataset are indicated below the dot. Dot size indicates significance of differential expression in the single-cell CRISPR screen and color indicates significance of differential expression in patients.

#### SUPPLEMENTAL TABLE CONTENTS

**Table S1. Multiplexed scRNA-seq of interacting NK cells and blood cancer cells**

(A) Cell line characteristics
(B) Multiplexed scRNA-seq samples, hashtag oligonucleotides, and summary statistics
(C) Differentially expressed genes between clusters of expanded NK cells
(D) Differentially expressed genes between clusters of PBMC NK cells
(E) Differentially expressed genes between untreated and expanded NK-cell treated cancer cells (all cell lines combined)
(F) Differentially expressed genes between untreated and PBMC NK-cell treated cancer cells (all cell lines combined)
(G) Differentially expressed genes between untreated and expanded NK-cell treated cancer cells (cell lines individually)
(H) Differentially expressed genes between untreated and PBMC NK-cell treated cancer cells (cell lines individually)
(I) Comparison of expanded NK-cell treated cancer cell DEG and genes correlated with cytolytic score in DLBCL (Reddy et al.)
(J) Comparison of expanded NK-cell treated cancer cell DEG and genes correlated with cytolytic score in AML (TCGA)
(K) Interactions between expanded NK cell clusters and cancer cells

**Table S2. PRISM-based NK cell sensitivity and its molecular correlates across blood cancer cell lines**

(A) Cell line annotations, PRISM barcodes, and PRISM-based NK cell sensitivity as AUC values
(B) PRISM-based NK cell sensitivity as percent viability across effector-to-target ratios
(C) Multi-omic correlations of PRISM-based sensitivity to NK cells

**Table S3. Genome-scale CRISPR screens of NK cell resistance**

(A) CRISPR screen technical specifications
(B) GeCKO v2 screen samples
(C) Sequencing primers
(D) K562 LOF
(E) MOLM14 LOF
(F) SUDHL4 LOF
(G) NALM6 LOF
(H) MM1S LOF
(I) LP1 LOF
(J) KMS11 LOF (KHYG1)
(K) MM1S GOF
(L) LP1 GOF (KHYG1)
(M) KMS11 GOF
(N) KMS11 GOF (KHYG1)
(O) Single gene validation sgRNAs
(P) Focused library sgRNAs

**Table S4. Genetic alterations and transcriptional regulation of CRISPR screen hits in blood cancer patients**

(A) CRISPR screen hit mutations in blood cancers
(B) Correlations of CRISPR screen hits to CNAs and methylation
(C) Differential expression of CRISPR screen hits in blood cancer compared to solid tumor cell lines

**Table S5. Integration of PRISM and CRISPR screens and patient data**

(A) Correlations of expression of concordant CRISPR/PRISM hits with PRISM-based sensitivity to NK cells across cancer types
(B) Multi-omic correlations of PRISM-based sensitivity to NK cells in AML
(C) Multi-omic correlations of PRISM-based sensitivity to NK cells in T-ALL
(D) Multi-omic correlations of PRISM-based sensitivity to NK cells in BCL

(E) Multi-omic correlations of PRISM-based sensitivity to NK cells in MM
(F) Multi-omic correlations of PRISM-based sensitivity to NK cells in B-ALL
(G) PRISM correlations in MM (CoMMpass)
(H) PRISM correlations in T-ALL (Liu et al.)
(D) PRISM correlations in DLBCL (Chapuy et al.)

**Table S6. Single-cell transcriptomics CRISPR screens of NK cell sensitivity regulators**

(A) CROP-seq sgRNA sequences and summary statistics
(B) Differentially expressed genes with each perturbation compared to control (K562)
(C) Differentially expressed genes with each perturbation compared to control (SUDHL4)
(D) Differentially expressed genes with each perturbation compared to control (NALM6)
(E) Differentially expressed genes with each perturbation compared to control (MM1S)
(F) Differentially expressed genes with each perturbation compared to control (LP1)
(G) Differentially expressed genes with each perturbation compared to control (MM1S GOF)
(H) Concordantly differentially expressed genes with CROP-seq perturbations and patients with mutations in the same genes
